# Oncogenic EGFR rewires STING-TBK1 immune machinery by tyrosine phosphorylation to license DNA damage tolerance

**DOI:** 10.1101/2025.10.20.683412

**Authors:** Qin Shen, Chen Mei, Yidan Chen, Qian Zhang, Qingzhe Wu, Xinyuan Yu, Fei Zhang, Shengduo Liu, Chen Chen, Cunqi Ye, Qi Zhang, Xin-Hua Feng, Hai Song, Tingbo Liang, Penghong Song, Bing Xia, Pinglong Xu

**Affiliations:** MOE Laboratory of Biosystems Homeostasis and Protection, Zhejiang Provincial Key Laboratory for Cancer Molecular Cell Biology, Life Sciences Institute, Zhejiang University, Hangzhou, 310058, China; Institute of Intelligent Medicine, ZJU-Hangzhou Global Scientific and Technological Innovation Center, Hangzhou, 310058, China; Department of Hepatobiliary and Pancreatic Surgery and Zhejiang Provincial Key Laboratory of Pancreatic Disease, The First Affiliated Hospital, University School of Medicine, Zhejiang University, Hangzhou, 310058, China; Department of Thoracic Cancer, Affiliated Hangzhou Cancer Hospital, Westlake University, Hangzhou, 310002, China; Cancer Center, Zhejiang University, Hangzhou, 310058, China

**Author notes:** Address for correspondence: Department of Hepatobiliary and Pancreatic Surgery, The First Affiliated Hospital, School of Medicine, Zhejiang University, 79 Qingchun Road, Hangzhou 310003, China, Bing Xia, Department of Thoracic Oncology, Hangzhou Cancer Hospital, Westlake University, Hangzhou, 310002, China, Pinglong Xu, Life Sciences Institute, Zhejiang University, 866 Yuhangtang Road, Nano Building, Room A555, Hangzhou, 310058, China. These authors contributed equally (Qin Shen, Chen Mei, Yidan Chen).

**Keywords:** EGFR, Hotspot Mutation, cGAS-STING, TBK1, Chemoresistance, EGFR-TKIs, tyrosine phosphorylation, Targeted therapy, Patient-Derived Organoids, Non-small cell lung cancer (NSCLC), Innate Immunity, Oncogenic signaling, Immune evasion, DNA damage repair, STING signalosome, Cisplatin, TBK1 inhibitor, Cancer Immunology

## Abstract

EGFR hotspot mutations (mEGFR), including primary L858R, exon 19 deletion, and secondary T790M, are pivotal oncogenic drivers in human non-small cell lung cancer (NSCLC). Meanwhile, NSCLC resistance to third-generation tyrosine kinase inhibitors (TKIs) is a major clinical challenge and remains mechanistically unresolved. Here, we uncover a previously unrecognized immunological mechanism whereby mEGFR exploits cGAS-STING innate immune signaling, conventionally regarded as tumor-suppressive, to sustain oncogenic signaling and therapeutic resistance. Mechanistically, mutant EGFR kinase aberrantly incorporates into STING signalosomes, directly phosphorylating STING (Y245/Y314) and TBK1 (Y577/Y677), stabilizing and hyperactivating TBK1 proteins, and establishing an unexpected and kinase loop critical for DNA damage repair. Disruption of this mEGFR-STING-TBK1 axis, genetically or pharmacologically, profoundly sensitized resistant patient-derived NSCLC organoids to chemotherapy. Combining TBK1 inhibition with cisplatin notably eradicated mEGFR-driven tumors in spontaneous and immunocompetent NSCLC murine models and patient-derived organoids. Our findings suggest a new function of cGAS-STING in the DNA damage repair program, its paradoxical exploitation by oncogenic driver mutations, and an innate immune therapeutic vulnerability in NSCLC.

## Introduction

Hyperactivation, overexpression, or genetic alterations in epidermal growth factor receptor (EGFR) contribute to the development and progression of an array of cancers, including breast cancer, lung cancer, colorectal and esophageal cancers, head and neck cancers, and glioblastoma^1,2^. Notably, hotspot mutations in the EGFR gene, especially the L858R mutation in exon 21 and deletions in exon 19, constitute 85-90% of all EGFR mutations observed in non-small cell lung cancer (NSCLC)^3,4^. These mutations lead to constitutive activation of the EGFR tyrosine kinase^5,6^, sustaining persistent oncogenic signaling through Ras-ERK, PLCγ1, PI3K-Akt, and STAT pathways^7^, thereby promoting cancer cell proliferation and survival while conferring a significant growth advantage. NSCLC driven by EGFR mutations initially responds to first- and second-generation TKIs, including erlotinib^8^, gefitinib^9^, and afatinib^10^. However, resistance frequently arises through secondary mutations^11,12^, particularly the T790M variant, and the activation of alternative signaling pathways^13^. Third-generation EGFR TKIs, such as osimertinib^14,15^, can overcome T790M mutation, but acquired resistance continues to emerge^16^, highlighting the urgent need to develop novel therapeutic strategies. These TKIs provide a lifeline for patients with specific resistance mutations, while immunotherapy and chemotherapy are standard subsequent treatment options when resistance occurs^17,18^. Cisplatin, a platinum-based chemotherapeutic agent that induces DNA damage and subsequent cell death^19^, has been used for decades to treat various solid tumors, including NSCLC^20^. However, the clinical efficacy of cisplatin is often compromised by the emergence of drug resistance^21^, occurring through numerous mechanisms, such as increased DNA repair, drug efflux, alterations in cell survival signals, epigenetic regulation, and evasion of apoptosis^20,22^. Therefore, elucidating the underlying mechanisms of next-generation TKIs and chemotherapeutic agents is paramount to designing more effective therapeutic strategies.

Understanding the molecular mechanisms involved in mEGFR-driven NSCLC, such as the interplays between mEGFR and innate immune signaling pathways, is crucial for developing treatment strategies against TKI and chemotherapy resistance. cGAS-STING (cyclic GMP–AMP synthase–stimulator of interferon genes) innate immune signaling triggers an array of physiological and pathological processes, including cytokine production, autophagy, and senescence, and is critical in host defense^23,24^, autoimmune diseases^25^, neurodegenerative diseases^26–28^, organ fibrosis^29,30^, and antitumor immunity^31–33^. Upon detecting mislocalized dsDNA, cGAS employs ATP and GTP molecules to synthesize 2’3’-cGAMP, the second messenger to activate STING proteins. Upon activation, STING transitions from the endoplasmic reticulum to the Golgi apparatus, and this migration enables its interaction with TBK1 (TANK-binding kinase 1) and triggers the activation of IRF3 (Interferon regulatory factor 3) and NF-κB (Nuclear factor kappa-light-chain-enhancer of activated B cells). These transcription factors transcribe type I interferons and other immune response genes, critical for cellular defense mechanisms and homeostasis^34^.

cGAS-STING signaling is traditionally considered a major initiator of antitumor immunity. The intratumoral application of STING agonists provokes robust immune- mediated tumor eradication in murine models, emphasizing the therapeutic potential of targeted STING activation in cancer cells and immune cells ^35,36^. Contrary to prior expectations, our research reveals an unexpected mechanism by which EGFR hotspot mutations hijack the STING-TBK1 pathway, repurposing it from a tumor-suppressive immune sensor into a pro-survival driver of chemoresistance. These mutations amplify STING signaling through direct interaction and enzymatic stabilization of TBK1, activating pro-survival pathways and reinforcing DNA repair machinery in mutant cells. We restore chemosensitivity by targeting this aberrant EGFR-STING-TBK1 axis, unveiling a previously unrecognized therapeutic vulnerability and a novel framework for overcoming TKI and chemotherapy resistance in EGFR-mutant NSCLC.

## Results

### EGFR hotspot mutations kinase-dependently facilitate cGAS-STING signaling

EGFR attracts considerable attention due to its hyperactivation in various cancer pathologies, particularly in NSCLC^37^. Roles of NSCLC-related EGFR hotspot mutations in cGAS-STING signaling were investigated. Intriguingly, we found that NSCLC-related EGFR hotspot mutants enhanced, instead of suppressing, IRF3 transactivation stimulated by STING but not MAVS (mitochondrial antiviral signaling protein) (sFig. 1A-B). The EGFR L858R/T790M, the most widely distributed mEGFR upon NSCLC targeted therapy, displayed a potent effect in amplifying STING signaling in a dose-dependent manner (sFig. 1C). In agreement with these observations, STING-TBK1-IRF3 signaling triggered by the STING agonists diABZI was substantially attenuated when mEGFR activity was abrogated by the third-generation tyrosine kinase inhibitor WZ4002 and osimertinib in mEGFR-driven NSCLC cells H1975 (with L858R/T790M mutation), as evidenced by severely compromised levels of phospho-TBK1 and IRF3 (Fig.1A). Conversely, H1299 NSCLC cells (with wild-type EGFR) were unresponsive to EGFR inhibitors and failed to influence STING signaling upon EGFR inhibition (Fig.1A). STING signaling stimulated by its natural agonist 2’3’-cGAMP, rather dsRNA sensing MAVS signaling, was impeded by mEGFR inhibitors (sFig. 1D). Given that STING and TBK1 clustering are key indicators of STING signaling activation, we measured these event in mEGFR-driven H1975 cells treated with diABZI. A more diffuse clustering pattern of endogenous STING and TBK1 proteins was observed upon mEGFR inhibition in H1975 cells (Fig. 1B), suggesting an essential role of mEGFR activity in STING signaling activation in these mEGFR-driven NSCLC cells. The effect of mEGFR inhibition on STING signaling was extended to other NSCLCs with EGFR mutations, such as in HCC827 cells harboring an EGFR delE746-A750 mutation (Fig. 1C). Consistent with these observations, depleting mEGFR using distinct shRNAs similarly curbed cGAS-STING signaling activated by synthetic B-DNA analogs in H1975 (Fig. 1D) or diABZI in HCC827 NSCLC cells(Fig. 1E). To validate this intriguing observation, we deleted endogenous mEGFR in H1975 cells by CRISPR/Cas9-mediated genome editing, which revealing severely impeded cGAS-STING signaling (Fig. 1F). Consistently, ectopic expression of mEGFR in H1299 cells significantly enhanced diABZI-induced STING signaling (Fig. 1G). We observed that mEGFR inhibitor WZ4002 substantially impeded mRNA levels of interferons and ISGs in diABZI-stimulated H1975 and HCC827 cells (Fig. 1H, sFig. 1E). These unexpected observations suggest that EGFR hotspot mutations, but not wild-type EGFR, is indispensable for cGAS-STING activation in NSCLC cells.

**Figure 1.**
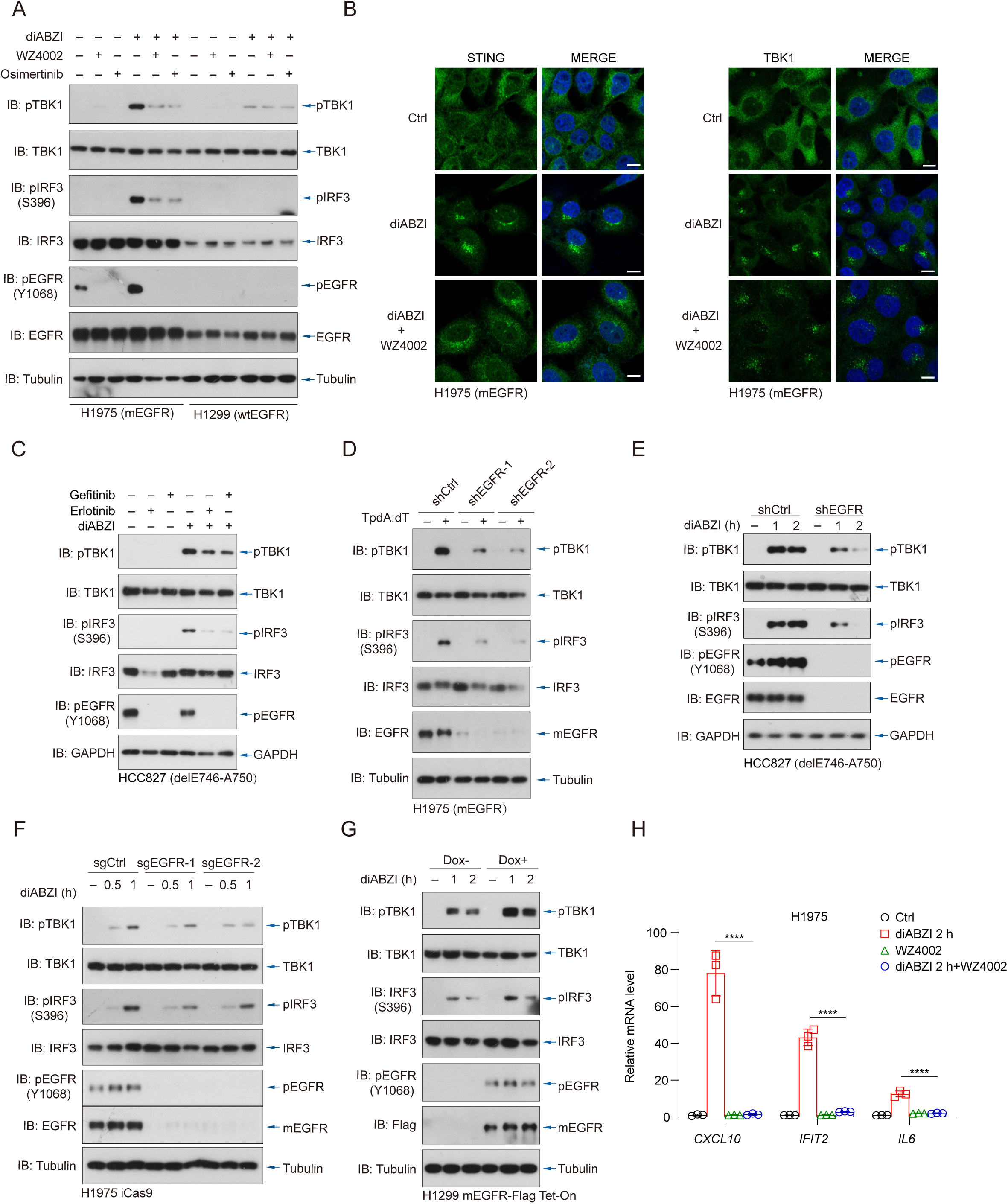
EGFR hotspot mutations kinase-dependently facilitate cGAS-STING signaling. **(A)** H1975 (EGFR L858R/T790M) and H1299 (wild-type EGFR) cells were treated with the STING agonist diABZI (10 μM, 1 h), with or without third-generation EGFR inhibitors WZ4002 or osimertinib (10 μM, 6 h). Immunoblotting was performed to assess the phosphorylation and total protein levels of TBK1, IRF3, and EGFR. **(B)** Immunofluorescence staining of STING (left) and TBK1 (right) in H1975 cells following diABZI stimulation (1 h) ± WZ4002 (6 h), revealing subcellular clustering. Scale bars, 10 μm. **(C)** HCC827 cells (EGFR delE746-A750) were treated with diABZI (1 h) with or without erlotinib or gefitinib (10 μM). Immunoblotting was used to analyze TBK1, IRF3, and EGFR activation. **(D)** H1975 cells expressing control or EGFR-targeting shRNAs were stimulated with poly(dA:dT) for 4 hours, followed by immunoblotting analysis. **(E)** EGFR knockdown HCC827 cells were treated with diABZI for the indicated time points, and phosphorylation of TBK1 and IRF3 was analyzed. **(F)** H1975 iCas9 cells transduced with control or EGFR-targeting sgRNAs were treated with diABZI and subjected to immunoblotting. **(G)** H1299 EGFR L858R/T790M Tet-On cells were treated with or without doxycycline (Dox) for 48 h, followed by diABZI stimulation for 1 or 2 hours. TBK1 and IRF3 activation was evaluated by immunoblotting. **(H)** H1975 cells were treated with diABZI (2 h), WZ4002, or both. mRNA levels of the interferon-stimulated genes *CXCL10*, *IFIT2*, and *IL6* were measured by quantitative RT-PCR. Data are presented as mean ± SEM (n = 3) and shown relative to untreated control cells. Statistical analysis was performed using one-way ANOVA with Bonferroni correction. ****P < 0.0001.

### mEGFR integrates into STING signalosomes

To investigate the mechanisms by which mEGFR potentiates STING signaling, we generated DLD1 cells with inducible expression of Flag-tagged EGFR wild-type, L858R/T790M, and delE746-A750 (sFig. 2A). Upon diABZI stimulation, we observed significant colocalization between mEGFR and STING/TBK1 clusters, indicating a substantial participation of mEGFR in STING signalosomes (Fig. 2A, sFig. 2B). In contrast, wild-type EGFR displayed no colocalization with STING/TBK1 puncta. This association between mEGFR and STING/TBK1 was abrogated in the presence of the mEGFR inhibitor WZ4002 (Fig. 2A, sFig. 2B), suggesting that the recruitment of mEGFR to the STING signalosome is kinase-dependent. Similarly, endogenous EGFR L858R/T790M proteins in H1975 cells exhibited robust colocalization with STING/TBK1 puncta following diABZI treatment, an interaction that was entirely abolished by mEGFR inhibitor WZ4002 (Fig. 2B-C). Immunofluorescence imaging also revealed differences in subcellular distribution for mEGFR and wtEGFR, as mEGFR localized partially at the Golgi apparatus and endosome, while wtEGFR mainly anchored to the plasma membrane (sFig. 2C).

**Figure 2.**
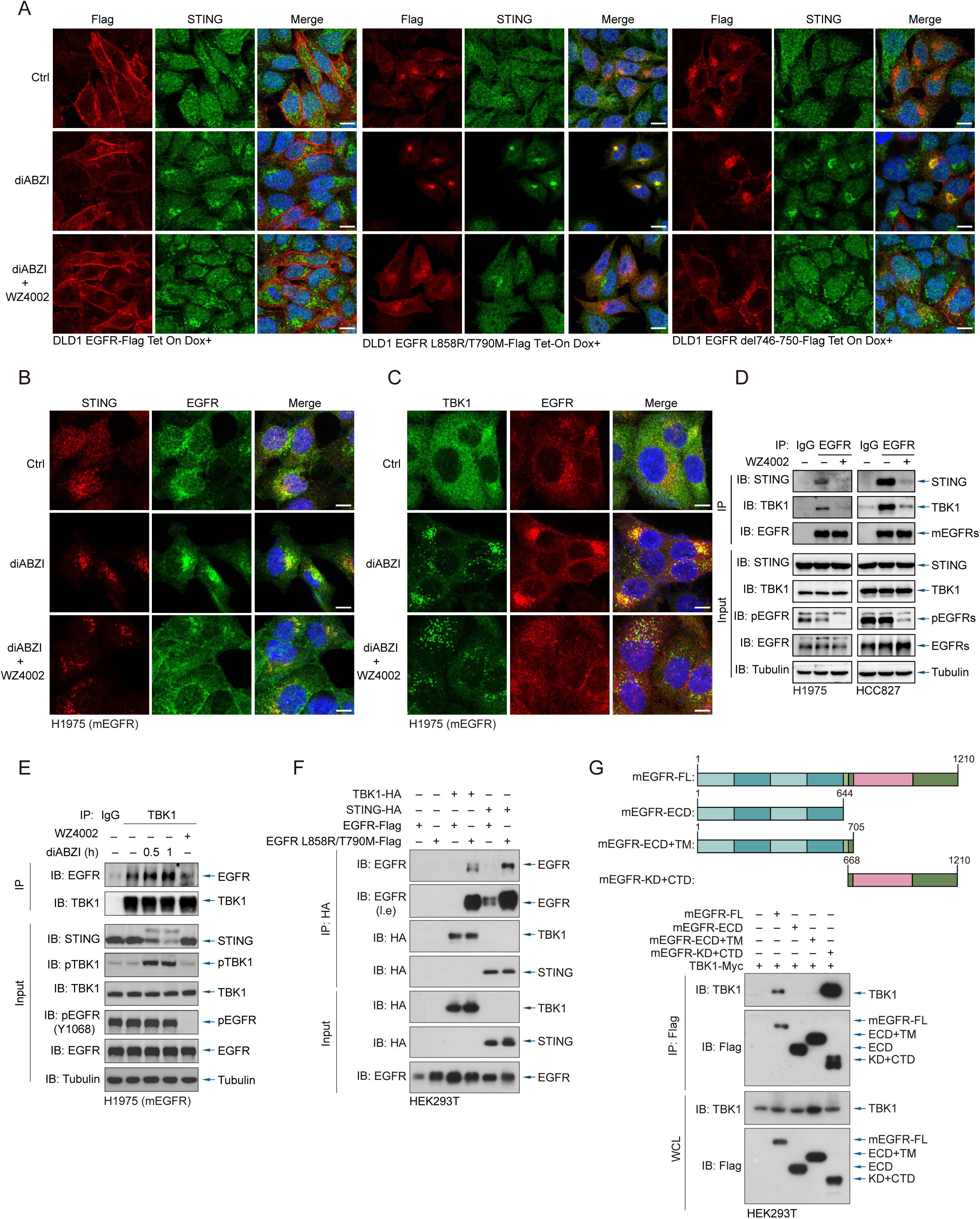
mEGFR, but not wild-type EGFR, integrates into STING signalosomes. **(A)** DLD1 Tet-On cells stably expressing Flag-tagged wild-type EGFR, EGFR L858R/T790M, or EGFR delE746-A750 were induced with doxycycline (Dox) and treated with diABZI (1 h) in the presence or absence of WZ4002 (6 h). Cells were fixed and stained with antibodies against Flag (red) and STING (green), and nuclei were counterstained with DAPI (blue). Representative immunofluorescence images show the subcellular localization and colocalization of EGFR variants with STING. Scale bars, 10 μm. **(B-C)** H1975 cells were treated with diABZI with or without WZ4002 and stained with antibodies against endogenous EGFR and STING (B), or EGFR and TBK1 (C). Scale bars, 10 μm. **(D)** H1975 and HCC827 cells were exposed to WZ4002 (10 μM, 6 h) or vehicle. Cell lysates were immunoprecipitated using anti-EGFR or isotype-matched IgG control antibodies and probed for STING, TBK1, pEGFR, and EGFR by immunoblotting. **(E)** H1975 cells were treated with diABZI (0.5 or 1 h), with or without WZ4002. Lysates were immunoprecipitated with anti-TBK1 or IgG control, and analyzed for EGFR association by immunoblotting. **(F)** HEK293T cells were transfected with Flag-tagged wild-type EGFR or EGFR L858R/T790M, HA-tagged TBK1 or STING, or combinations. After 24 hours, lysates were immunoprecipitated with anti-HA antibody and analyzed by immunoblotting with anti-EGFR and anti-HA antibodies. **(G)** Top: Schematic of EGFR truncation constructs used for interaction mapping, including full-length (FL), extracellular domain only (ECD), ECD plus transmembrane domain (ECD+TM), and kinase domain with C-terminal tail (KD+CTD). Bottom: HEK293T cells were co-transfected with Flag-tagged EGFR L858R/T790M truncation constructs and Myc-tagged TBK1. Anti-Flag immunoprecipitants were analyzed by immunoblotting for Flag and TBK1.

Coimmunoprecipitation experiments further validated these observations in mEGFR-expressing H1975 and HCC827 cells, where mEGFR robustly interacted with endogenous STING and TBK1 proteins in a kinase-dependent manner (Fig. 2D). Additionally, endogenous complexes of mEGFR with TBK1 in H1975 cells were somewhat enhanced following diABZI stimulation (Fig. 2E). By contrast, wild-type EGFR barely interacted with STING and TBK1, even when ectopically expressed (Fig. 2F).

Intriguingly, mEGFR, unlike wild-type EGFR, markedly enhanced the associations of TBK1 with both its upstream adaptor STING and downstream effector IRF3, indicating the role of mEGFR in STING signalosome assembly (sFig. 2D-E). We analyzed their association using protein truncation mutants to determine whether the interaction between mEGFR and STING signalosomes is membrane trafficking-based or direct protein-protein interactions. Kinase- and C-terminal domains of mEGFR were robust and sufficient to interact with TBK1 (Fig. 2G). These findings consistently indicate that mEGFR, but not wtEGFR, is an integral component of classical STING-TBK1-IRF3 signalosomes.

### Tyrosine phosphorylation of STING and TBK1 by mEGFR solidifies STING clustering

To elucidate mechanisms by which mEGFR integrates STING machinery and modulates STING signaling, we first utilized an array of inhibitors targeting directly mEGFR or its downstream signaling components, including AKT, Erk1/2, and STAT3/5^38^. Inhibitors targeting mEGFR kinase activity impeded STING-induced phosphorylation of TBK1 and IRF3, while inhibitors acting on AKT, Erk1/2, and STAT3/5 did not (Fig. 3A). Phos-tag PAGE analyses further confirmed a substantial increase in TBK1 phosphorylation levels (phospho-S172) upon mEGFR co-expression, rather than wild-type EGFR (Fig. 3B). Noticeably, we detected a robust phospho-tyrosine signal (pY100) on both TBK1 (Fig. 3C) and STING proteins (Fig. 3D) following mEGFR expression, but not in IKKε or by wild-type EGFR. Tyrosine phosphorylation of both endogenous STING and TBK1 proteins was seen in H1975 NSCLC cells harboring mEGFR but not in H1299 NSCLC cells harboring wtEGFR (Fig. 3E). mEGFR kinase inhibitor WZ4002 significantly diminished these phosphorylations (Fig. 3E), illustrating the presence of tyrosine phosphorylation of endogenous STING and TBK1 proteins in mEGFR-driven NSCLC.

**Figure 3.**
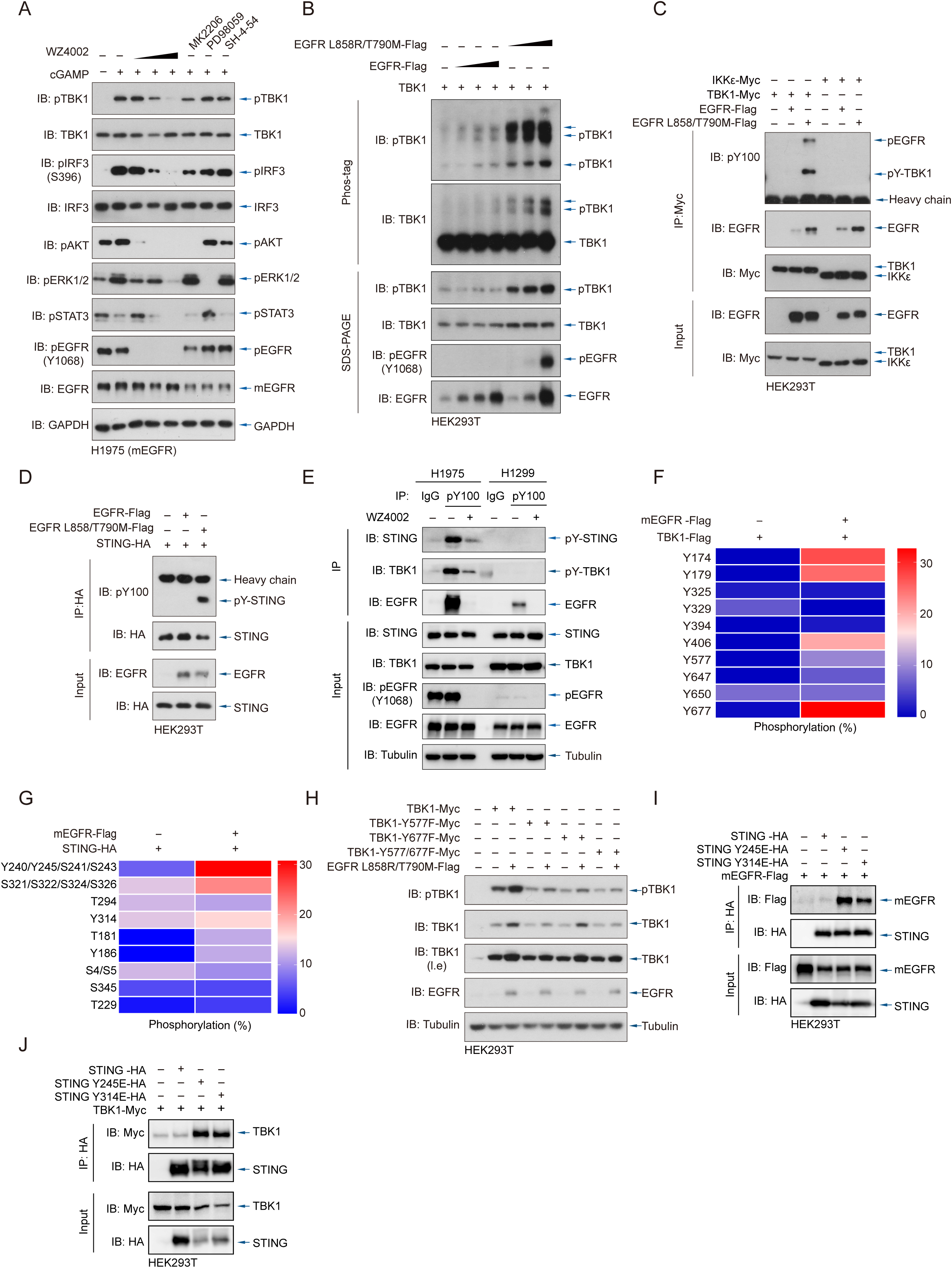
mEGFR tyrosine phosphorylates and facilitate STING and TBK1 activation. **(A)** H1975 cells were pretreated with increasing concentrations of EGFR inhibitor WZ4002 (0.1, 1, or 10 μM) or with specific pathway inhibitors MK2206 (AKT inhibitor), PD98059 (MEK1 inhibitor), or SH-4-54 (STAT3 inhibitor), each at 10 μM for 6 h, followed by stimulation with cGAMP (1 h). Whole cell lysates were analyzed by immunoblotting. **(B)** HEK293T cells were transfected with Flag-tagged wild-type EGFR or EGFR L858R/T790M, along with Myc-tagged TBK1. Cell lysates were analyzed by conventional SDS-PAGE or Phos-tag gel electrophoresis to assess TBK1 phosphorylation. **(C)** HEK293T cells were co-transfected with Flag-tagged wild-type or mutant EGFR, along with TBK1-Myc or IKKε-Myc. Lysates were immunoprecipitated with anti-Myc and analyzed by immunoblotting with anti-phosphotyrosine (pY100) and anti-EGFR to detect tyrosine phosphorylation of TBK1 and IKKε. **(D)** HEK293T cells were co-transfected with HA-tagged STING with either wild-type or mutant EGFR. HA immunoprecipitants were probed with pY100 and STING antibodies to assess STING tyrosine phosphorylation. **(E)** H1975 and H1299 cells were treated with or without WZ4002 (10 μM, 6 h). Lysates were immunoprecipitated with pY100 antibody, and analyzed for STING, TBK1, and EGFR by immunoblotting. **(F)** Mass spectrometry was performed on TBK1 immunoprecipitants from HEK293T cells co-expressing TBK1 and EGFR L858R/T790M. The heatmap displays relative tyrosine phosphorylation levels at specific TBK1 residues. **(G)** HEK293T cells were transfected with HA-tagged STING, with or without co-expression of EGFR L858R/T790M. Mass spectrometry heatmaps indicate relative phosphorylation at tyrosine and serine/threonine sites in STING. **(H)** Wild-type or mutant TBK1 constructs (Y577F, Y677F, or Y577F/Y677F) were co-expressed with EGFR L858R/T790M in HEK293T cells. TBK1 phosphorylation was reduced in single mutants and diminished in the double mutant. YF: tyrosine to phenylalanine **(I)** HA-tagged wild-type STING or phosphomimetic mutants (Y245E, Y314E) were co-expressed with Flag-tagged EGFR L858R/T790M. HA immunoprecipitants were blotted for EGFR and STING to assess protein interaction. Increased binding was observed between mutant EGFR and phosphomimetic STING mutants. YE: tyrosine to glutamate. **(J)** HA-tagged wild-type or phosphomimetic STING mutants were co-expressed with Myc-tagged TBK1. HA immunoprecipitation followed by immunoblotting revealed enhanced binding of TBK1 to STING Y245E/Y314E compared to wild-type STING.

Next, mass spectrometry analyses identified several tyrosine residues of TBK1 (Fig. 3F) and STING (Fig. 3G) as potential phosphorylation targets by mEGFR. We then generated a series of TBK1 and STING tyrosine-to-phenylalanine mutants guided by mass spectrometry data. mEGFR coexpression revealed a notable reduction in pY100 signals in TBK1 Y577F, Y677F, or Y577F/Y677F mutants (sFig. 3A), including the Y577 residue that forms a cation–π interaction with R375, a key interaction necessary for the integration of TBK1 into the STING signalosomes^39^. By contrast, mEGFR-enhanced TBK1 activation was abolished in the Y577F/Y677F mutant (Fig. 3H), proposing Y577 and Y677 residues as key sites for mEGFR-mediated TBK1 regulation. Meanwhile, immunoblotting following coimmunoprecipitations indicated that STING Y245 and Y314 residues were the main sites responsible for mEGFR-mediated phosphorylation (sFig. 3B). Noticeably, the phosphomimetic mutants of STING Y245E and Y314E exhibited substantial affinity to both mEGFR (Fig. 3I) and TBK1 (Fig. 3J), when compared to wild-type STING, suggesting a tyrosine phosphorylation-based recruitment of mEGFR and TBK1 into STING signalosomes. These data suggest that mEGFR tyrosine phosphorylates STING and TBK1, solidifying mEGFR recruitment and potentiating STING signalosome assembly.

### TBK1 phosphorylation at Y577/Y677 stabilizes activated TBK1

Besides enhancing STING signalosome assembly that facilitates TBK1 activation, does mEGFR also regulate the magnitude or duration of TBK1 activation? TBK1 is degraded by the ubiquitin-proteasomal system^40,41^. We observed an increase in TBK1 protein levels upon mEGFR cotransfection, and inducible expression of mEGFR diminished the diABZI-induced degradation of TBK1 (Fig. 4A, sFig. 4A). Blocking proteasome function by MG132 inhibited diABZI and cGAMP-triggered degradation of endogenous TBK1 in H1975 cells (Fig. 4B, sFig.4B), suggesting a role of mEGFR in stabilizing activated TBK1 upon STING activation. In the presence of cycloheximide, which blocked de novo protein synthesis, mEGFR significantly delayed TBK1 degradation (Fig. 4C). Furthermore, mEGFR, rather than wild-type EGFR, notably inhibited the K48-Ub modification of transfected TBK1 proteins in HEK293T cells (Fig. 4D). Endogenous TBK1 proteins in NSCLC cells (Fig. 4E). Noticeably, phosphomimetic TBK1 mutant (Y577E/Y677E, 2YE) exhibited a markedly extended half-life under cycloheximide chase assay (Fig. 4F). Consistently, a minimal level of K48-linked ubiquitination on the phosphomimetic TBK1 mutant was detected when compared to WT or 2YF TBK1 mutant. However, their K63-linked ubiquitination remained comparable (Fig. 4G-H). Lentivirus-delivered reconstitution of mutant TBK1 in TBK1-deleted H1299 cells also indicated that STING agonist-induced K48-Ub modification on TBK1 was abrogated in the case of TBK1-2YE (Fig. 4I), with higher protein levels than wild-type TBK1 (Fig. 4J). These findings collectively suggest that mEGFR-mediated phosphorylation of TBK1 at Y577/Y677 protected it from K48-Ub modification and proteasomal degradation, thus sustaining STING-TBK1 signaling.

**Figure 4.**
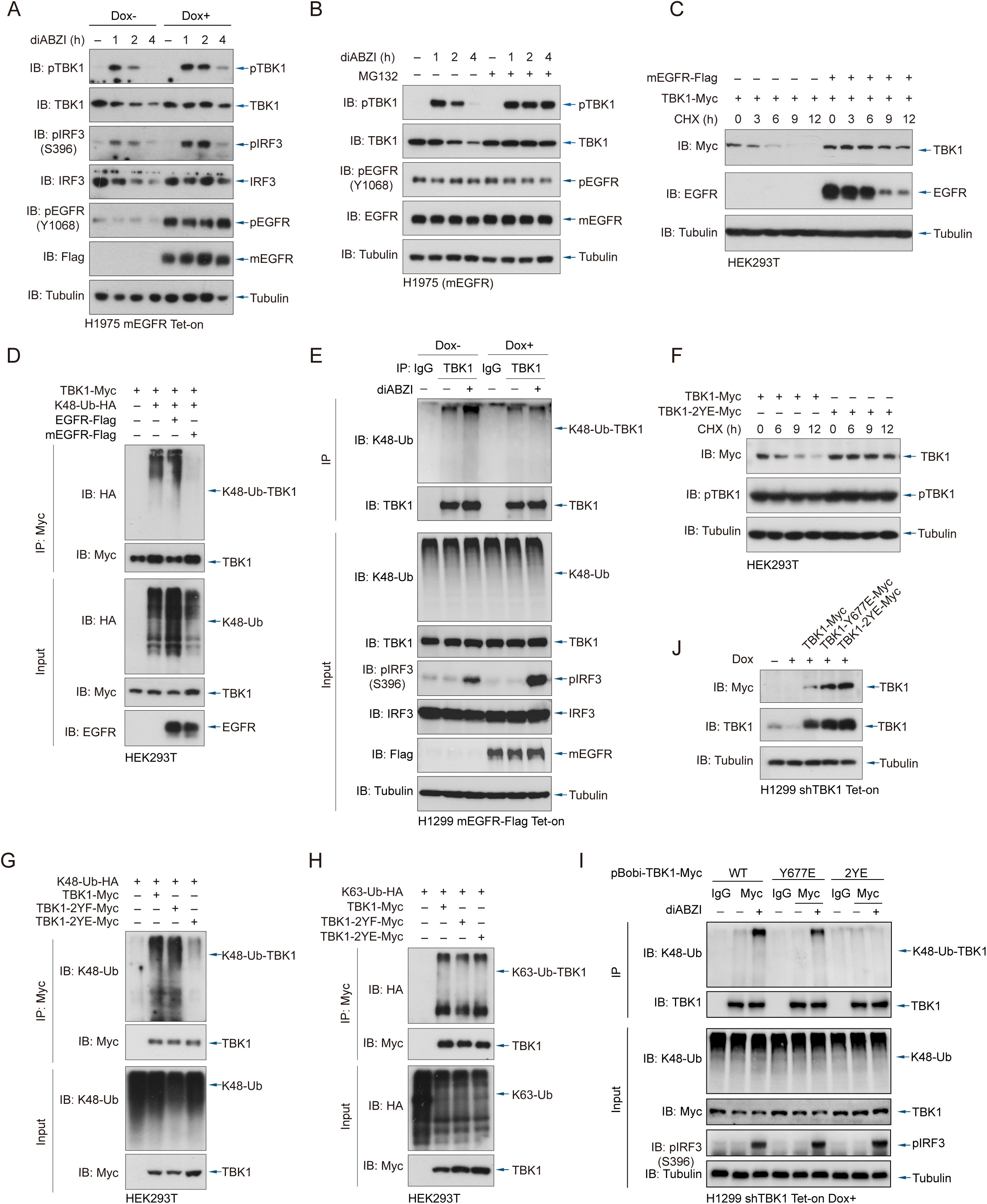
mEGFR-mediated phosphorylation hyperactivates and stabilizes TBK1. **(A)** H1975 cells stably expressing doxycycline-inducible EGFR L858R/T790M were treated with diABZI for the indicated time points with or without doxycycline pre-treatment (48 h). TBK1 protein levels were analyzed by immunoblotting to assess degradation dynamics. **(B)** H1975 cells were treated with diABZI for the indicated time points in the presence or absence of MG132 (10 μM, 6 h) to assess the effect of proteasomal inhibition on TBK1 degradation. **(C)** HEK293T cells were co-transfected with TBK1-Myc with or without Flag-tagged EGFR L858R/T790M. Cells were treated with cycloheximide (CHX) for the indicated time points to assess TBK1 protein stability by immunoblotting. **(D)** HEK293T cells were co-transfected with TBK1-Myc, HA-tagged K48-Ubiquitin (K48-Ub), and either wild-type or mutant EGFR. TBK1 ubiquitination was examined by immunoprecipitation with anti-Myc followed by anti-K48-Ub immunoblotting. **(E)** H1299 cells stably expressing doxycycline-inducible EGFR L858R/T790M were treated with diABZI (1h) with or without doxycycline induction. Endogenous TBK1 was immunoprecipitated and analyzed for K48-linked ubiquitination. **(F)** HEK293T cells were transfected with either wild-type TBK1 or TBK1 phosphomimetic mutant (Y577E/Y677E, designated as TBK1-2YE). TBK1 protein stability was assessed by CHX chase assays and immunoblotting. **(G)** TBK1-Myc (wild-type, 2YF, or 2YE) was co-expressed with K48-Ub in HEK293T cells. TBK1 was immunoprecipitated and analyzed for K48-linked ubiquitination by immunoblotting to evaluate the role of phosphorylation in ubiquitin modification. **(H)** K63-linked ubiquitination of TBK1 was examined by co-transfecting K63-Ub-HA with TBK1 variants (WT, 2YF, or 2YE) in HEK293T cells. Immunoprecipitated TBK1 was analyzed for K63-Ub by immunoblotting. **(I)** H1299 cells with doxycycline-inducible knockdown of endogenous TBK1 were reconstituted with shRNA-resistant TBK1-Myc constructs (WT, Y677E, or 2YE). Cells were treated with diABZI (1 h), followed by immunoprecipitation using anti-Myc and analysis of K48-linked ubiquitination. **(J)** The steady-state expression levels of TBK1 variants (WT, Y677E, 2YE) were analyzed by immunoblotting in H1299 Tet-On shTBK1 cells after 72 hours of doxycycline-induced knockdown of endogenous TBK1.

### STING and TBK1 are both indispensable for mEGFR-driven NSCLC survival

Hyperactivation of EGFR is a significant driver of cancer development, promoting tumorigenesis through uncontrolled proliferation, survival, and metastasis^42^. However, our findings above suggest an unexpected role of mEGFR in promoting cGAS-STING signaling, an innate immune system armed for antitumor immunity. Does mEGFR- potentiated cGAS-STING signaling contribute to protumorigenic functions? Noticeably, we found that doxycycline-induced shRNA depletion of either TBK1 or STING substantially inhibited H1975 growth to a level comparable to mEGFR depletion (Fig. 5A), while TBK1 depletion significantly reduced cell viability in mEGFR-driven NSCLC cells harboring L858R/T790M (Fig. 5B) and delA746-E750 (sFig. 5A). Moreover, an elevated level of cleaved-PARP in H1975 cells following TBK1 or STING depletion was detected (Fig. 5C), and Annexin-FITC/PI assays revealed apparent apoptosis of mEGFR- driven H1975 NSCLC cells following doxycycline-induced TBK1 depletion (Fig. 5D), suggesting mEGFR-activated STING signalosomes in preventing NSCLC apoptotic death. Additionally, TBK1 depletion potently impeded colony formation of mEGFR-driven NSCLC cells (Fig. 5E), while barely impacting colony formation in H1299 cells harboring wild-type EGFR, suggesting a unique dependency and vulnerability of mEGFR-driven cells on TBK1 (Fig. 5F). Consistently, the reconstitution of the TBK1 2YE mutant into H1975 cells, combined with a silent mutation of the shRNA recognition site, wholly rescued the colony formation defect caused by TBK1 depletion (Fig. 5G), suggesting an essential role of mEGFR-directed TBK1 stability in survival regulation.

**Figure 5.**
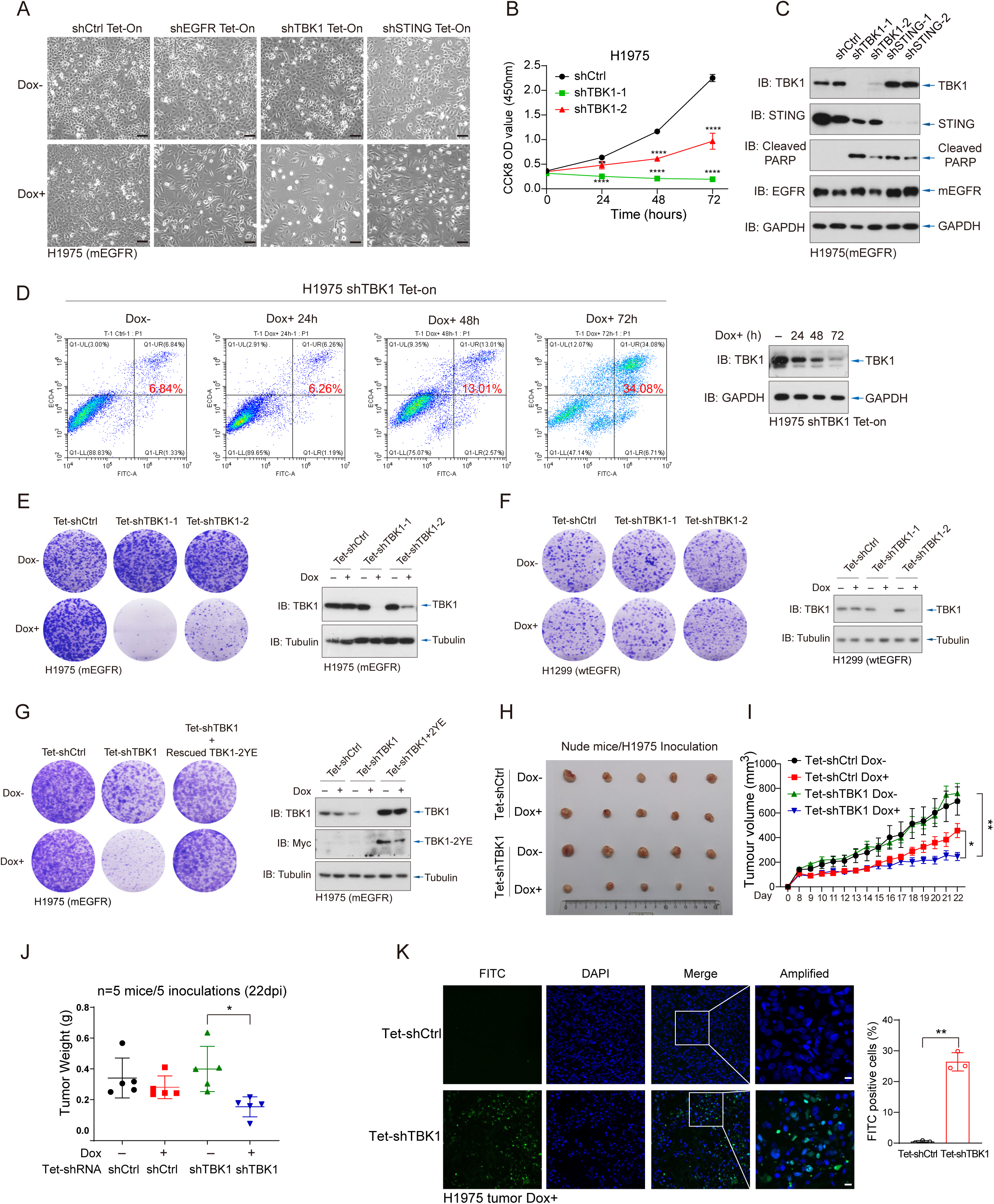
STING and TBK1 are indispensable for mEGFR-driven NSCLC survival. **(A)** Brightfield images of H1975 cells expressing inducible shRNAs targeting EGFR, TBK1, STING, or a non-targeting control (shCtrl). Cells were treated with Dox or left untreated for 72 hours. Scale bar, 100 μm. **(B)** Cell viability of H1975 cells expressing two independent shRNAs targeting TBK1 (shTBK1) or shCtrl was assessed at the indicated time points using a CCK-8 assay. **(C)** Immunoblotting confirmed knockdown efficiency of TBK1 and STING in H1975 cells. Apoptosis was evaluated by detection of cleaved PARP following TBK1 or STING depletion. **(D)** Apoptosis in H1975 cells expressing Tet-On shTBK1 was analyzed at 24, 48, and 72 hours of Dox induction using Annexin V-FITC/PI staining and flow cytometry. Percentages of Annexin V-positive cells are indicated. Corresponding TBK1 levels were validated by immunoblotting. **(E-F)** Colony formation assays of H1975 (E) and H1299 (F) cells expressing inducible shCtrl or two independent shTBK1 constructs, with or without Dox treatment. Knockdown efficiency of TBK1 was confirmed by immunoblotting (right panels). **(G)** Colony formation assays of H1975 cells with TBK1 knockdown and reconstitution of an shRNA-resistant TBK1-2YE. Expression of TBK1 and TBK1-2YE was validated by immunoblotting. **(H)** Representative tumor images from nude mice subcutaneously inoculated with H1975 cells expressing Tet-On shCtrl or shTBK1. Mice were fed with or without doxycycline-containing (1mg/mL) water for 22 days post-inoculation. **(I)** Tumor growth curves of H1975 xenografts with or without Dox-induced TBK1 depletion (n = 5 per group). Tumor volumes were measured at the indicated time points. **(J)** Tumor weights of H1975 xenografts collected at 22 days post-inoculation (dpi) from mice treated with or without doxycycline (n = 5). **(K)** FITC-labeled TUNEL staining of tumor sections (n = 3) from H1975 xenografts expressing Tet-On shCtrl or shTBK1. DAPI (blue) stains nuclei, and FITC (green) indicates apoptotic cells. Scale bars, 10 μM. Quantification of FITC- positive cells is shown on the right. Data in B and I-K represent mean ± SEM. Statistical analysis by ANOVA with Bonferroni correction; *, P<0.05, **, P<0.01, and ***, P<0.001.

Next, we employed an in vivo immune-independent model to validate the proposed role of STING signalosomes in mEGFR-driven NSCLC viability. H1975 cells with inducible shCtrl or shTBK1 shRNA were subcutaneously injected into immune-incompetent mice, and doxycycline-induced TBK1 depletion in H1975 cells significantly attenuated tumor growth and tumor weight (Fig. 5H-J, sFig. 5B-C). Apoptosis level of mEGFR-driven H1975 tumors upon TBK1 depletion was also verified by TUNEL staining, an effect not observed in shCtrl tumors (Fig. 5K). Collectively, these findings establish STING signalosomes as an indispensable factor, through TBK1 activation, for the maintenance and progression of mEGFR-driven NSCLC cells.

### TBK1 facilitates DNA damage repair programs to confer cisplatin resistance

Next, we performed transcriptional profiling in H1975 cells to assess the impact of TBK1 depletion on the transcription landscape. RNAseq analyses showed substantial alterations in gene expression upon TBK1 depletion (Fig. 6A). KEGG enrichment analysis indicated significant involvement of pathways related to p53 signaling, cell cycle regulation, and DNA replication (sFig. 6A). Gene Set Enrichment Analysis (GSEA) further highlighted the activation of TNF signaling, IL-17 signaling, and other pathways associated with inflammation and immune responses. Significant downregulation of DNA repair pathways was observed following TBK1 depletion, including base excision repair, homologous recombination repair, and mismatch repair (Fig. 6B-C). Consistent with GSEA findings, we observed a notable increase in apoptotic markers in TBK1- deficient cells in response to TNFα treatment (sFig. 6B). Additionally, colony formation assays showed a substantial reduction in the number of colonies when the TBK1 inhibitor GSK8612 or TBK1 depletion was combined with TNFα treatment (sFig. 6C). These observations suggest that TBK1 plays an essential role in facilitating NSCLC cell survival by regulating survival programs such as DNA damage repair. The mechanisms underlying TBK1-promoted DNA damage repair are not fully understood and are probably involved in PARP9^43^, ATM^44^, ISG15^45^, and deoxyribonucleotide metabolism^46^, which are not pursued in the current study.

**Figure 6.**
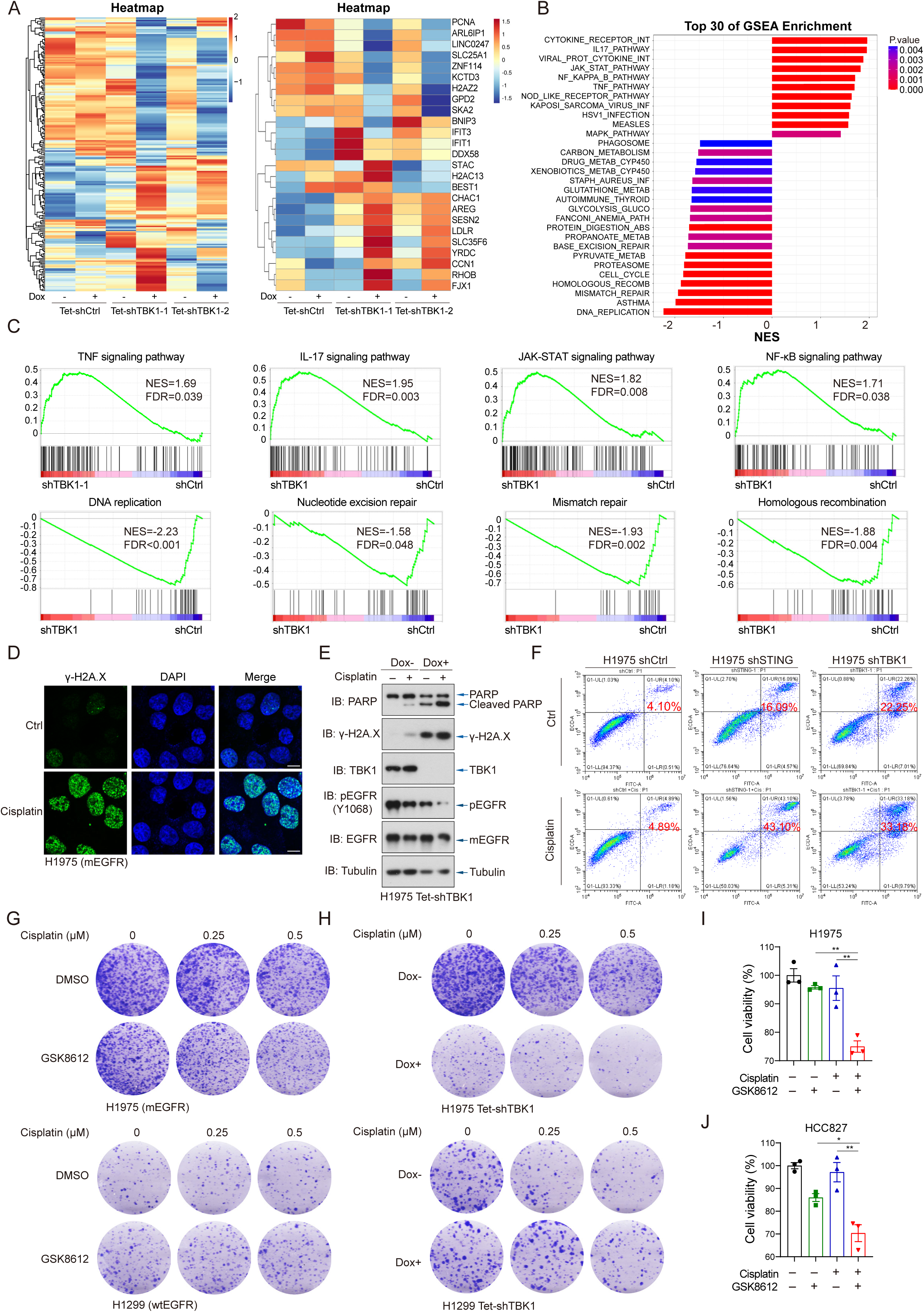
TBK1 facilitates DNA damage repair programs to survive mEGFR-driven NSCLC. **(A)** Heatmap of transcriptomic profiles from H1975 cells expressing doxycycline-inducible TBK1 shRNAs (Tet-shTBK1-1 and Tet-shTBK1-2) or control shRNA (Tet-shCtrl), with or without Dox induction. Left: hierarchical clustering of differentially expressed genes. Right: top upregulated and downregulated genes. Color scale denotes relative expression levels from low (blue) to high (red). **(B)** Gene Set Enrichment Analysis (GSEA) revealed significant suppression of DNA repair pathways upon TBK1 depletion in H1975 cells. **(C)** Representative GSEA plots demonstrating enrichment of inflammatory signaling and impairment of DNA damage repair pathways in TBK1-depleted cells. NES, normalized enrichment score; FDR, false discovery rate **(D)** Representative immunofluorescence images of γ-H2A.X (green), a marker of DNA double-strand breaks, in H1975 cells treated with cisplatin (8 µM, 24 h). Scale bars, 10 µm. **(E)** Immunoblotting of H1975 Tet-shTBK1 cells treated with cisplatin (8 µM, 24 h), with or without Dox, showed increased levels of cleaved PARP and γ-H2A.X, indicating enhanced apoptosis and impaired DNA repair upon TBK1 depletion. **(F)** Flow cytometry analysis of Annexin V/PI-stained H1975 cells expressing shCtrl, shSTING, or shTBK1, treated with cisplatin (1 µM, 48 h). Depletion of STING or TBK1 significantly enhanced cisplatin-induced apoptosis. **(G)** Colony formation assays in H1975 and H1299 cells treated with cisplatin (doses as indicated), with or without the TBK1 inhibitor GSK8612 (5 µM). **(H)** Colony formation assays in Tet-shTBK1 H1975 and H1299 cells treated with cisplatin, with or without Dox-induced TBK1 knockdown. **(I-J)** CCK-8 assays measuring viability of H1975 (I) and HCC827 (J) cells treated with cisplatin (0.4 µM) and/or GSK8612 (5 µM). Combination treatment significantly reduced cell viability compared to single-agent treatments. Data in I-J are presented as mean ± SEM. Statistical analysis by ANOVA with Bonferroni correction; *P < 0.05; **P < 0.01; ***P < 0.001.

Cisplatin is a well-established clinical chemotherapeutic agent, particularly in cases without targetable mutations, ineligible for immunotherapy, and targeted therapy resistance^47^, and cisplatin requires efficient DNA repair mechanisms for resistance^20^. We confirmed that cisplatin treatment significantly increased γ-H2A.X foci formation in H1975 cells (Fig. 6D). Noticeably, we found that cisplatin robustly activated cGAS-STING-TBK1-IRF3 signaling in mEGFR-driven H1975 and HCC827 NSCLC cells (sFig. 6D-F), as well as STING-dependent expression of interferon-stimulated genes (ISGs) (sFig. 6G). Coimmunoprecipitation assays indicated that cisplatin facilitated the formation of EGFR-STING-TBK1 tripartite complexes, thus potentially linking mEGFR to STING signalosome activation and drug resistance. Both γ-H2A.X and cleaved PARP were significantly increased in TBK1-depleted NSCLC cells upon cisplatin treatment (Fig. 6E), and STING- or TBK1-deficient NSCLC cells exhibited significantly increased levels of apoptosis upon a low dose of cisplatin treatment (Fig. 6F).

Moreover, combinational treatment of cisplatin and TBK1 inhibition substantially and synergistically attenuated colony formation in H1975 cells (Fig. 6G). By contrast, this combinatorial strategy did not affect H1299 cells harboring wild-type EGFR (Fig. 6G). Likewise, TBK1 deficiency by dox-induced shRNA rendered H1975, but not H1299, more sensitive to cisplatin-induced cytotoxicity (Fig. 6H). In both H1975 and HCC827 cells, treatment with cisplatin or GSK8612 alone led to modest reductions in cell viability, while their combination significantly reduced NSCLC viability (Fig. 6I-J). Dose-response curves further confirmed that cisplatin IC_50_ in H1975 and HCC827 cells was substantially reduced upon TBK1 inhibition (sFig. 6I-J) or STING depletion (sFig. 6K-L). These data suggest that targeting STING signaling in mEGFR-driven NSCLC impairs DNA damage repair and sensitizes tumors to chemotherapeutic agents.

### Combining cisplatin and TBK1i eradicates TKI-resistant NSCLC PDOs and mEGFR- driven spontaneous NSCLC tumors

H1975 tumors engrafted in NSG mice indicated the TBK1 inhibitor GSK8612 substantially intensified the effect of cisplatin, achieving complete elimination in half of the tumors upon a month-long regimen (sFig. 7A-B), and quantitative assessments of both tumor volume and weight established the superior efficacy of the drug combination over monotherapy (sFig. 7C-D). TUNEL staining illustrated a marked elevation in apoptosis ratio within the tumor tissue (sFig. 7E), suggesting an impressive antitumor effect of cisplatin and GSK8612 combinational therapy in an artificial immune-deficient environment.

Next, we explored a synergistic treatment of GSK8612 and cisplatin in patient-derived organoids (PDO) and spontaneous murine NSCLC models. Organoids derived from EGFR-mutant patients who had developed resistance to 3^rd^-generation TKIs and one with wild-type EGFR were established and evaluated for synergistic drug effects. Cell viability assays indicated that the combination of cisplatin and TBK1 inhibitor GSK8612 significantly reduced the viability of mEGFR PDOs but not PDO bearing wild-type EGFR (Fig. 7A). A substantial synergistic effect between cisplatin and GSK8612 was confirmed by Combination Index (CI) analysis (sFig. 7F). Dose-response curves confirmed the enhanced cisplatin sensitivity in mEGFR-PDOs by GSK8612 (Fig. 7B), suggesting a specific synthetic lethal interaction between TBK1 inhibition and cisplatin in EGFR-mutant NSCLC. The real-time apoptotic response was measured in PDOs by caspase green apoptosis probe (CGAP) assays, monitoring a rapid and substantially increased caspase activity in organoids treated with the combination of cisplatin and GSK8612, compared to the monotherapy (Fig. 7C-D, sFig. 7G), suggesting a synergistic effect in eradicating mEGFR-driven and TKI-resistant NSCLC carcinoma by cisplatin and GSK8612.

**Figure 7.**
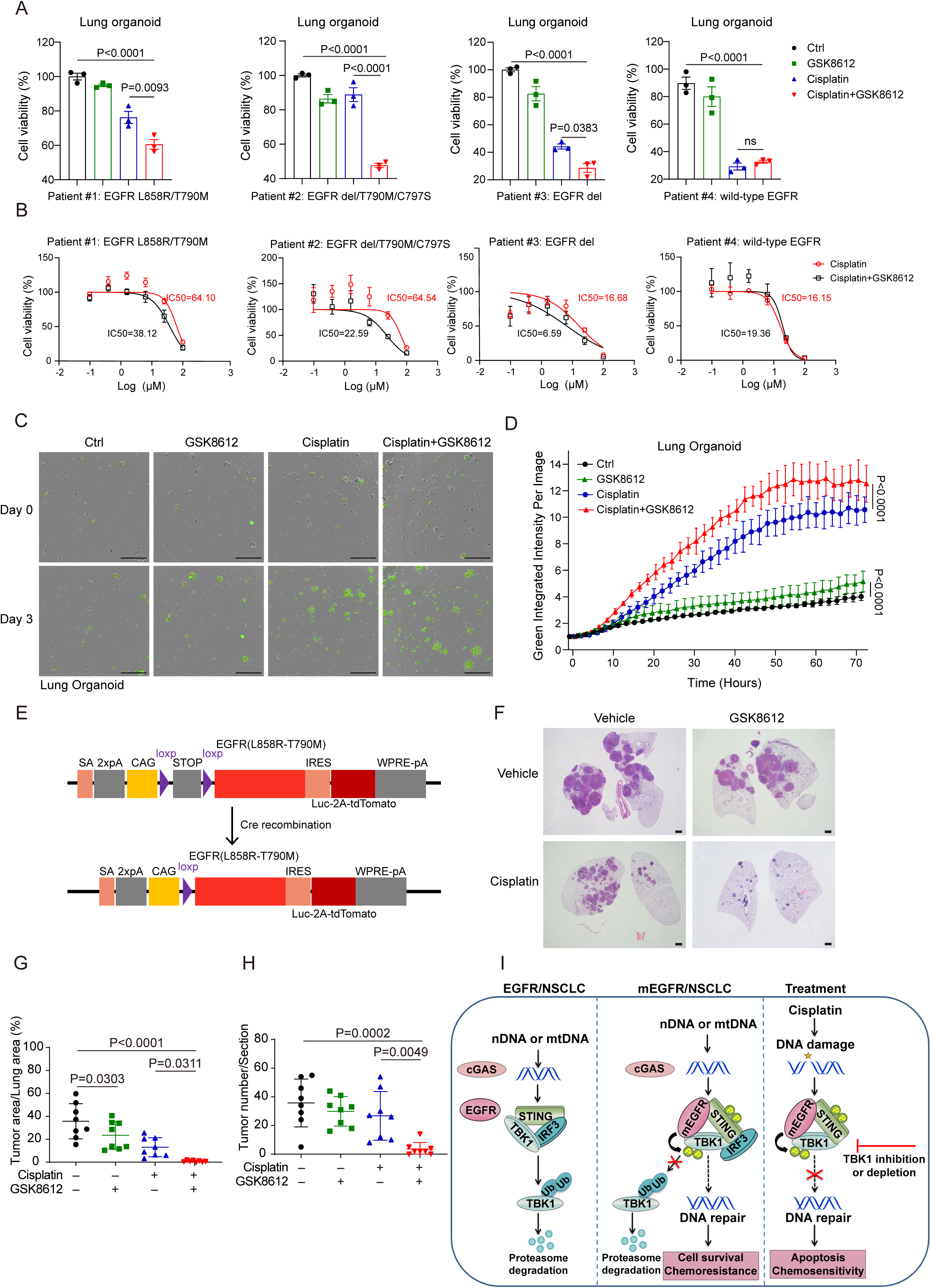
Combining cisplatin and TBK1i eradicates mEGFR-driven NSCLC PDOs and spontaneous tumors. **(A)** Cell viability assays using CellTiter-Lumi™ were performed on patient-derived lung organoids (PDOs) from NSCLC patients bearing EGFR mutations (L858R/T790M, exon19 del/T790M, or exon19 del) or wild-type EGFR. Combination treatment with cisplatin (25 μM) and TBK1 inhibitor GSK8612 (5 μM) significantly reduced cell viability in PDOs with mutant EGFR, but not in wild-type EGFR organoids. ns, not significant **(B)** Dose–response curves for cisplatin alone or combined with GSK8612 were generated for each PDO. The combination treatment markedly reduced IC₅₀ values in EGFR-mutant organoids (Patients #1–3), while wild-type EGFR organoids (Patient #4) showed no appreciable sensitization. **(C)** Representative fluorescence images of PDOs (Patient #1) treated with indicated conditions for 72 h, stained using Caspase-3/7 Green ReadyProbes™ to detect apoptotic cells (green). Scale bars, 500 μM. **(D)** Quantification of caspase-3/7 fluorescence intensity over time in PDOs from Patient #1. Combined treatment with cisplatin and GSK8612 induced significantly higher apoptosis than either monotherapy. Data are presented as mean ± SEM (n=3). Statistical analysis was performed by two-way ANOVA with Bonferroni correction. **(E)** Schematic of the Cre-inducible genetically engineered mouse model expressing EGFR L858R/T790M and luciferase-tdTomato reporter, allowing conditional activation of mEGFR-driven lung tumors upon Cre recombination. **(F)** Representative hematoxylin and eosin (H&E) staining of lung sections from mice treated with vehicle, GSK8612, cisplatin, or their combination. Scale bars, 100 μM. **(G–H)** Quantification of tumor burden in murine lungs. Combined treatment with cisplatin and GSK8612 significantly reduced tumor area relative to total lung area (**G**) and decreased tumor number per lung section (**H**), compared with either monotherapy. Data are shown as mean ± SEM (n=8). **(I)** Proposed working model: mutant EGFR (mEGFR) engages and tyrosine-phosphorylates STING and TBK1, integrating into STING signalosomes and sustaining TBK1 activation, which promotes chemoresistance and tumor survival. Therapeutically, co-targeting TBK1 with chemotherapy synergistically suppresses EGFR-driven NSCLC.

mEGFR-driven lung carcinogenesis in an immunocompetent and genetically engineered murine model was next employed, from which Rosa26^mEGFR^ transgenic mice developed spontaneous tumors upon Adeno-Cre virus instillation (Fig. 7E). Since most advanced NSCLC patients often developed p53 loss, these mice were subjected to CRISPR-LentiV2-Cre-mediated p53 knockout to expedite tumorigenesis and followed by a month-long combinational therapy regimen (sFig. 7H). These immunocompetent mice developed mEGFR-driven carcinomas, and immunohistochemistry revealed robust intratumoral pEGFR signaling (sFig. 7I). The combined therapy of cisplatin and GSK8612, surpassing cisplatin or GSK8612 monotherapy, markedly attenuated tumor progression, as evidenced by hematoxylin and eosin (HE) tissue staining (Fig. 7F) and the measurement of tumor size and tumor numbers (Fig. 7G-H), suggesting a critical role of TBK1 oncogenic signaling, rather than classic STING-mediated antitumor immunity, in controlling mEGFR-driven NSCLC. Therefore, exploiting this STING-TBK1 vulnerability of mEGFR-driven NSCLC can offer a promising strategy against chemoresistance to eliminate tumor burden in PDOs and preclinical NSCLC models.

## Discussion

Our study reveals a novel biological event by which oncogenic EGFR mutations in NSCLC exploit innate immune signaling for survival, thereby challenging the conventional view of cGAS-STING as a tumor suppressor. We demonstrate that these activating EGFR mutations integrate physically into STING signalosomes and form a self-reinforcing loop strengthened by EGFR-mediated STING phosphorylation. mEGFR noticeably modifies TBK1, inhibiting its degradation and extending its activation magnitude and duration. Crucially, TBK1 becomes indispensable for the survival of EGFR-mutant NSCLC cells and orchestrates a prosurvival signaling network involving improved DNA damage repair (Fig. 7I). Disrupting this aberrant EGFR-STING-TBK1 axis restores sensitivity to chemotherapy in both NSCLC cells and patient-derived NSCLC organoids. Therefore, this study fundamentally shifts our understanding of cGAS-STING signaling and offers a potential therapeutic vulnerability for overcoming chemoresistance in EGFR-mutant NSCLC.

### EGFR hotspot mutations integrate and facilitate cGAS-STING signaling

While EGFR mutations are well-established drivers of NSCLC progression and resistance, their interactions with cGAS-STING signaling remain largely unexplored. Emerging evidence highlights a probable dual role of cGAS-STING signaling in antitumor immunity^48,49^. In breast cancer, chromosomal instability drives metastasis in a STING-dependent manner, highlighting its dual role in cancer progression^50^. STING activation can also sustain tumor survival by inducing IL-6-dependent pro-survival signaling in chromosomally unstable cancers^51^ and an inflammatory microenvironment promoting low-grade inflammation and recruiting myeloid-derived suppressor cells (MDSCs)^52,53^. In tumors with low antigenicity, STING also fosters tumor growth through indoleamine 2,3-dioxygenase (IDO)-mediated immune suppression^54^. These complexities highlight the necessity for a comprehensive understanding of the specific implications of the cGAS-STING pathway in distinct cancer contexts.

Our study uncovered an interplay between mEGFR and cGAS-STING in NSCLC cells. Specifically, mEGFR variants, including L858R/T790M, were found to integrate into STING signalosomes and facilitate a robust activation of STING-TBK1-IRF3 signaling, as evidenced by increased phosphorylation of STING and TBK1 and elevated expression of interferon-stimulated genes (ISGs). This interaction was exclusive to mEGFR, as wild-type EGFR neither exhibited the same integration within STING signalosomes nor induced comparable STING activation^32^ (sFig. 1A-B). These findings suggest that mEGFR evolves and uniquely hijacks STING machinery, potentially driving oncogenic immune signaling in NSCLC.

A few mechanisms explain the selective recruitment of mEGFR into STING signalosomes. First, mEGFR-mediated tyrosine phosphorylation of STING creates a docking signal, amplifying their association and generating a self-reinforcing feedback loop within the signalosome (sFig. 3C-D). Second, the elevated endocytosis rate of mEGFR compared to wild-type EGFR^55^ might increase its co-localization with STING in endosomal compartments. Immunofluorescence analyses further revealed distinct subcellular localization patterns of mEGFR and wild-type EGFR, even under basal conditions where spatial proximity facilitates interaction (Fig. 2A-B). Additionally, direct physical interactions between mEGFR and STING, potentially strengthened by adaptor proteins, could further stabilize their co-localization and enhance signaling. The ability of mEGFR, but not wild-type EGFR, to enhance STING signaling likely stems from its increased kinase activity. mEGFR harbors activating mutations, such as L858R and T790M, which potentiate its catalytic function and modify its structural conformation^56^, potentially exposing interaction domains critical for binding to STING and TBK1. Our data indicate that the kinase domain and C-terminal region of mEGFR are essential for interacting with STING and TBK1, suggesting a direct role in modulating innate immune signaling. Therefore, the affinity distinction between wild-type and EGFR hotspot mutations to STING constitutes the molecular basis for the acquired ability of mEGFR in driving NSCLC tumorigenesis.

### mEGFR utilizes TBK1 kinase to facilitate DNA damage repair for chemoresistance

In NSCLC cells expressing mEGFR, TBK1 is markedly activated and stabilized within STING signalosomes, evidenced by enhanced phosphorylation of TBK1 at its activation sites and a prolonged half-life in mEGFR-driven cells (Fig. 4A). TBK1 can activate PI3K- AKT signaling under non-immune stresses^57,58^ and is involved in Kras-driven and VHL- deficient cancers^59,60^. Our findings reveal a previously unrecognized mechanism by which mEGFR initiates and extends the lifespan of activated TBK1. Specifically, mEGFR directly phosphorylates TBK1 at tyrosine residues Y577 and Y677, which inhibits K48- linked polyubiquitination, the key signal for TBK1 proteasomal degradation. This stabilization mechanism underlies the intricate regulatory network that mEGFR exploits to sustain TBK1 activity, reinforcing oncogenic immune signaling and promoting cancer cell survival. TBK1, the central kinase downstream of STING activation, has received substantial attention in cancer biology. Besides its recognized roles in antitumor immunity^61,62^, TBK1 can respond to cell stress and contribute to survival and proliferative signals^58,63,64^. As such, TBK1 prevents TNF-induced cell death by RIPK1 phosphorylation^65^ and serves as a synthetic lethal target in Kras-driven^66^ and VHL loss cancers^59^. TBK1 facilitates tumor survival by promoting the formation of an immunosuppressive environment, which could be a potential therapeutic target^67,68^. However, the potential role of TBK1 in chemoresistance and DNA damage repair remains poorly understood, particularly in mEGFR-driven NSCLC.

We found that TBK1 plays a pivotal role in the survival of mEGFR-driven NSCLC cells, functioning as a key regulator of pro-survival signaling networks. Upon activation, TBK1 can orchestrate multiple oncogenic pathways, including PI3K-AKT/mTORC1 and ERK MAPK/RIPK1, which drive cell growth, proliferation, and survival, ultimately fostering tumor progression and chemoresistance^57,69^. Our findings and previous studies have identified TBK1 as a central molecule within these pathways^68,70^, thus facilitating the viability of mEGFR-driven cancer cells under chemotherapeutic stress. Beyond its anticipated role in pro-survival signaling, our transcriptomic analysis surprisingly reveals that TBK1 suppression profoundly impairs DNA repair processes. TBK1 depletion significantly downregulates multiple DNA repair pathways, including base excision repair, homologous recombination, and mismatch repair (Fig. 6B-C). This reduction in DNA repair capacity likely sensitizes NSCLC cells to chemotherapy, particularly cisplatin, which induces DNA damage by forming DNA adducts and interstrand crosslinks^20^. Without efficient DNA repair, these lesions accumulate, pushing cells toward apoptosis^20^. Thus, TBK1 emerges as a dual facilitator of chemoresistance by activating pro-survival signaling and supporting DNA repair mechanisms that protect cancer cells from chemotherapy-induced damage. However, the mechanisms underlying TBK1-promoted DNA damage repair are not fully understood, and are probably involved in PARP9^43^, ATM^44^, ISG15^45^, and deoxyribonucleotide metabolism^46^.

Therefore, TBK1 is uniquely positioned at the crossroads of antitumor immunity and cancer cell survival. TBK1 is essential for activating immune responses against pathogens and foreign nucleic acids^71,72^. However, in mEGFR-driven NSCLC, TBK1 is hijacked to sustain cancer cell survival and drive resistance to chemotherapy, transforming it from an immune mediator into an oncogenic driver. This dual function presents TBK1 as a complex yet promising therapeutic target. Targeting TBK1 disrupts a key survival pathway in cancer cells and potentially restores antitumor immunity, as TBK1 has been identified as a critical immune evasion gene^73^. Our study validates the feasibility of this strategy, demonstrating that pharmacological inhibition of TBK1 sensitizes mEGFR-driven NSCLC cells to chemotherapy in both in vitro models and patient-derived organoids. Future research should focus on elucidating the upstream regulators and downstream effectors of TBK1 while developing more potent and selective TBK1 inhibitors for clinical application.

Therefore, this study unveils a significant conceptual leap by elucidating the unanticipated exploitation of cGAS-STING innate immune signaling in driving therapeutic resistance within EGFR-mutated NSCLC. Beyond addressing the immediate clinical importance of overcoming EGFR-TKI resistance, these findings expand our understanding of cancer biology by revealing a paradoxical mechanism wherein tumor cells evolutionally co-opt antitumor immune pathways to their advantage. This paradigm shift suggests that the subversion of innate immune pathways may be a common mechanism of tumor evolution and therapeutic resistance across diverse tumor types. While highlighting TBK1 as a promising therapeutic target in combination with chemotherapy for mEGFR-mutated NSCLC, these findings also raise critical questions concerning the failure of STING activation to elicit antitumor immunity in this context and the potential risks of STING agonist application in mEGFR-driven NSCLC.

## ACKNOWLEDGMENTS

This research was sponsored by the NSFC Projects (32321002, 32430028, and 31830052 to P.X., and 82271678 to Q.Z.), the National Key Research and Development Program of China (2021YFA1301401 to P.X.), Science and Technology Development Project of Hangzhou (20241029Y037 to Y.C.), Medical and Health Science and Technology Program Projects of Zhejiang Province (2025KY1113 to Y.C.), and NSFC-Zhejiang Province (Q22C079669 to S.L.).

## AUTHOR CONTRIBUTIONS

Q.S. and C.M. carried out most experiments. Y.C. and Q.Z. contributed to several experiments, Q.W., F.Z., S.L., C.C., C.Y., Q.Z., X.F., H.S., T.L., P.S., and B.X. helped with resources, data analyses, and discussions. P.X. and Q.S. conceived the study and experimental design. P.X. and Q.S. wrote the manuscript.

## METHOD DETAILS

### Expression plasmids, reagents, and antibodies

Expression plasmids encoding Flag-, Myc-, or HA-tagged wild-type or mutations, or the truncations of human MAVS, STING, IRF3-2SA, TBK1, IKKε, IRF3, K48-Ub, K63-Ub, and the reporters of IFNβ_Luc and 5xISRE_Luc have been previously described^74,75^. Open reading frames (ORFs) of human CALR, GOLGA1, and Rab5A were obtained from the Invitrogen ORF Lite Clone Collection cDNA library by PCR. Flag-human EGFR and NSCLC-related mutants were constructed on the pcDNA3.1 mammalian expression vector. Flag-tagged truncations of human mEGFR were constructed on the mammalian expression vector pRK5. Site-directed mutagenesis about TBK1 Y406F/E, TBK1 Y577F/E, TBK1 Y677F/E, TBK1 Y(406+677)F/E, TBK1 Y(577+677)F/E, STING-Y254F/E, STING-Y314F/E and STING-Y(245+314)F were generated by PCR-based cloning performed by a kit from Stratagene. EGFR, EGFR delE746-A750, and EGFR L858R/T790M were constructed on lentiviral vector pCMV, utilizing them to generate the Tet-On system in various cells. Myc-TBK1, Myc-TBK1-Y677E, and Myc-TBK1-Y(577+677)E were constructed on lentiviral pBobi, utilizing them to generate stable cell lines of H1299 and H1975 cells. All coding sequences were verified by DNA sequencing, and the detailed information for these constructions was provided in Supplementary Table 1. The pharmacological reagents Elrotinib (Selleck), Gefitinib (Selleck), WZ4002 (Selleck), Osimertinib (Selleck), GSK8612 (Selleck), Cisplatin (Selleck), MK2206 (Selleck), PD98059 (Selleck), SH-4-54 (Selleck), TRIzol (Yeasen), Doxycycline (Sangon Biotech), cycloheximide (CHX; Sigma), cGAMP (Invivogen), diABZI (Selleck), poly(I: C) (Invivogen), poly (dA:dT) (Invivogen), puromycin (Yeasen), G418 (Yeasen), and CMC-Na (Selleck) were purchased and used according to manual instructions.

Detailed information on antibodies applied in immunoblotting, immunoprecipitation, and immunofluorescence was provided in Supplemental Table 1. The monoclonal anti-STING (50494, 1:2000 dilution), anti-TBK1 (3504, 1:3,000 dilution), anti-TBK1 (38066, 1:500 dilution), anti-pTBK1(S172) (5483, 1:2,000 dilution), anti-IRF3 (4302, 1:2,000 dilution), anti-pIRF3 (S396) (4947, 1:2,000 dilution), anti-STAT1 (14994, 1:1,000 dilution), anti-pSTAT1 (S727) (8826, 1:1,000 dilution), anti-EGFR (4267; 1:3,000 dilution), anti-pEGFR (Y1068) (3777, 1:2,000 dilution), anti-pEGFR(Y845) (2231;1:2,000 dilution), anti-AKT1 (2938; 1:2,000 dilution), anti-pAKT1(S473) (4060; 1:2,000 dilution) anti-ERK1/2 (4695T; 1:2,000 dilution), anti-pERK1/2(T202 Y204) (4370T; 1:2,000 dilution), anti-pY100 (9411S; 1:1,000 dilution), anti-cleaved PARP (5625; 1:2,000 dilution), anti-K48-Ub (8081, 1:2,000 dilution), anti-K63-Ub (5621, 1:2,000 dilution), anti-Caspase-3 (9662S, 1:2,000 dilution), anti-PARP (9542S, 1:2,000 dilution), anti-γ-H2A.X (9718, 1:1,000 dilution), anti-GAPDH (5174S, 1:5,000 dilution), anti-Myc (2276S, 1:3,000 dilution), and anti-HA (3724S, 1:5,000 dilution) antibodies were purchased from Cell Signaling Technology. The anti-TBK1 (ab40676, 1:3,000 dilution), anti-IRF3 (ab68481, 1:3,000 dilution), anti-pIRF3 (S386) (ab76493, 1:2,000 dilution), anti-EGFR (ab52894, 1:3000 dilution), and anti-pY100(ab179530, 1:2000 dilution) were purchased from Abcam, and the anti-tubulin (T6199, 1:10,000 dilution), anti-β-actin (A5441, 1:10,000 dilution), anti-HA (H9658, 1:2000 dilution), and anti-Flag (M2) (F3165, 1:5,000 dilution) were purchased from Sigma. The anti-STING (19851-1- AP, 1:300 dilution) was purchased from Proteintech. Anti-GM130 (1:1,000 dilution) was purchased from BD Biosciences. The anti-STING (518172, 1:100 dilution), anti-EGFR (52894, 1:100 dilution), anti-Calnexin (23954, 1:50 dilution), anti-EEA1(53939, 1:100 dilution), anti-rabbit IgG, and anti-mouse IgG antibodies were purchased from Santa Cruz.

### Cell culture, transfection, and infection

H1299, HCC827, and H1975 cells were cultured in RPMI 1640 medium supplemented with 10% fetal bovine serum (FBS) at 37 °C in 5% CO2 (vol/vol). HEK293T and DLD1 cells were maintained in DMEM medium with 10% fetal bovine serum (FBS) at 37 °C in 5% CO2 (vol/vol). Inducible DLD1, H1299, and H1975 cells for expressing EGFR L858R/T790M were generated by infection of lentiviral vectors harboring the inducible Tet-On system and selected with the antibiotic G418 at a concentration of 1,500 μg ml^−1^ for DLD1 cells or 500 μg ml^−1^ for H1975 and H1299 cells for one week. Transfection reagents, Polyethyleneimine (PEI, Polysciences) or Lipofectamine RNAiMAX (Invitrogen), were used for the transfection of plasmids, poly(dA:dT), and poly(I: C). Digitonin (Sigma) at a concentration of 10 μg ml^−1^ was used to transport cGAMP into cells. Lentiviral infection was performed by directly applying a virus-containing medium into target cells, which was replaced by a fresh medium containing 10% FBS after 6 hours.

### Lung organoid culture

Tumor samples were obtained following ethical approval from the Ethics Committee of Hangzhou Cancer Hospital (Approval No. HZCH-2022-016, dated September 30, 2022), with four clinical cases: Patient #1, EGFR L858R/T790M and resistance to Almonertinib and Firmonertinib; Patient #2, EGFR del19+T790M+C797S and resistance to Osimertinib; Patient #3, del19 and resistance to Almonertinib; Patient #4, EGFR wild-type and treated with Pemetrexed, cisplatin, and Sintilimab. Tumor samples were stored at 4°C in preservation solution and processed within 48 hours. After removing excess tissue, samples were washed in DPBS-PS, minced to 0.5-1 mm³ in 200 µL of digestion solution I, and then digested in 4 mL of digestion solution I at 37°C for 1 hour. Digestion was stopped with 8 mL DPBS-PS and centrifugation at 1500 rpm for 5 minutes. The pellet was resuspended in 2 mL of digestion solution II, incubated at 37°C for 15 minutes, stopped with 4 mL DPBS-PS, and then centrifuged. Cells were filtered through a 100 µm strainer, recentrifuged at 1200 rpm, counted, and resuspended in Matrigel (5000-10,000 cells per 50 µL). Cell domes were plated in 24-well plates, solidified, and overlaid with 500 µL MasterAim® Lung Cancer Organoid Culture Medium. Organoids were cultured at 37°C in 5% CO₂.

### Luciferase reporter assay

HEK293T cells were transfected with the indicated reporters bearing an ORF coding for the Firefly luciferase alongside the pRL-Luc with the Renilla luciferase ORF as the internal control for transfection and other expression vectors specified in the results section. Briefly, after 12 h post-transfection, the cells were treated with the indicated inhibitors and lysed in a passive lysis buffer (Promega) 24 hours after transfection. The luciferase assays were performed using a dual luciferase assay kit (Promega), quantified with POLARstarOmega (BMG Labtech), and normalized to the internal Renilla-luciferase activity.

### Quantitative RT–qPCR assay

H1975 and HCC827 cells stimulated with diABZI and cisplatin were lysed, and total RNA was extracted using an RNAeasy extraction kit (Axygen). Complementary DNA was generated using a one-step iScript cDNA synthesis kit (Vazyme), and qRT-PCR was performed using EvaGreen qPCR master mixes (Abcam) and a CFX96 real-time PCR system(Bio-Rad). Relative quantification was expressed as 2^−ΔCt^, where C_t_ is the difference between the main C_t_ value of the triplicate of the sample and the C_t_ value of endogenous L19 mRNA. The following oligos were used: hIFIT1, 5’- TTGATGACGATGAAATGCCTGA -3’, 5’- CAGGTCACCAGACTCCTCAC -3’; hIFIT2, 5’-GACACGGTTAAAGTGTGGAGG -3’, 5’- TCCAGACGGTAGCTTGCTATT -3’; hCXCL10, 5’-GTGGCATTCAAGGAGTACCTC -3’, 5’- TGATGGCCTTCGATTCTGGATT -3’; hIFNβ1, 5’-ATGACCAACAAGTGTCTCCTCC -3’, 5’- GGAATCCAAGCAAGTTGTAGCTC-3’; hISG15, 5’-CGCAGATCACCCAGAAGATCG -3’, 5’- TTCGTCGCATTTGTCCACCA -3’; hIL6, 5’ - GTCAGGGGTGGTTATTGCAT - 3’, 5’ - AGTGAGGAACAAGCCAGAGC - 3’. All human primers used in the RT–qPCR assays were listed in Supplementary Table 1.

### Coimmunoprecipitation and immunoblotting

HEK293T, H1975, HCC827, and H1299 cells stimulated with cGAMP, diABZI poly (I: C), and poly (dA:dT) or transfected with the specified plasmids encoding Myc-, FLAG- or HA-tagged EGFR, mEGFRs, MAVS, TBK1, IRF3, or STING were lysed in a modified MLB lysis buffer (20 mM Tris-Cl, 200 mM NaCl, 10 mM NaF, 1 mM NaV2O4, 1% NP-40, 20 mM β-glycerophosphate, and protease inhibitor, pH 7.5)^76^. The cell lysates were then subjected to immunoprecipitation using antibodies against FLAG (Sigma, F3165-5MG; 1:200 dilution), Myc (CST, 2276S; 1:200 dilution) or HA (Sigma, H9658; 1:200 dilution) for transfected proteins or using anti-TBK1 (3504S, 1:200 dilution) or anti-pY100 (Abcam, 179530; 1:200 dilution) antibody for endogenous proteins. After three or four washes with cold MLB, the adsorbed proteins were resolved by SDS–PAGE (Bio-Rad) and subjected to immunoblotting with the indicated antibodies. Cell lysates were also analyzed using SDS-PAGE and immunoblotting to control the protein abundance.

### Immunofluorescence, microscopy, and FACS

To visualize the co-localization of the induced mEGFR with endogenous STING/TBK1, DLD1 cells expressing the inducible proteins specified in the Results section (including FLAG-tagged mEGFR or FLAG-tagged STING for 24 h) were treated either with cGAMP or WZ4002 for the indicated times, fixed in 4% paraformaldehyde, blocked in 2% BSA in PBS for 1 h, and incubated sequentially with primary antibodies—anti-FLAG (M2; Sigma, F3165; 1:300 dilution), anti-STING (Abcam, ab181125; 1:100 dilution), anti-EGFR, or anti-TBK1, and the Alexa-labelled secondary antibodies (Jackson Laboratories, 111-095-003, 115-095-003, 111-025-003, and 115-025-003; 1:500 dilution) with extensive washes. H1975 slides were then mounted with Vectorshield and stained with DAPI (Vector Laboratories). Immunofluorescence images were obtained and analyzed using a Nikon Eclipse Ti inverted microscope, Zeiss LSM710, Zeiss LSM880 confocal microscope, or OLYMPUS spinning disk confocal microscope. According to the manufacturer’s instructions, a BD FACSCalibur was used for FACS analysis of Annexin-FITC/PI-positive cells.

### shRNA-mediated RNA interference

The shRNA sequence was designed by GPP WEB PORTAL (https://portals.broadinstitute.org/gpp/public/ and listed in Supplementary Table 1. Oligonucleotides were cloned into pLKO.1 or Tet-pLKO.1 plasmid with the AgeI/EcoRI sites to generate mEGFR, TBK1, or STING-depleted cells. The infection process was similar to retroviral infection except that the lentiviral packaging plasmids PLP1, PLP2, and VSV-G were transfected into HEK293T cells for virus production. Media containing lentivirus were collected 48 hours after transfection. For two days, cells were selected in 5 µg/mL puromycin in RPMI 1640 medium.

### CRISPR/Cas9-mediated generation of knockout cells

To generate knockout cell lines, guide RNA sequences targeting human EGFR were designed using the CRISPRdb database (https://crisprdb.org/) and cloned into a lentiviral vector. HEK293T cells were transfected with the lentiviral packaging plasmids PLP1, PLP2, and VSV-G to produce the virus. The collected viral supernatants were used to transduce target cells, which were selected 24 hours post-infection using flow cytometry to isolate Cas9-GFP-positive cells. These cells were then treated with 5 µg/mL puromycin for 48 hours to ensure selection, and the efficiency of gene knockout was confirmed through Western blot analysis. All primers used in the CRISPR/Cas9 methods are also listed in Supplemental Table 1.

### Cell viability assay

To evaluate cell viability, H1975 and HCC827 cells transfected with control shRNA or shTBK1 were seeded at a density of 3,000 cells per well in 96-well plates. After an initial incubation to allow for cell attachment (37°C, 5% CO2), Cell viability was then measured at the indicated time points using the CCK8. 10 μL of CCK8 solution was added to each well for the assay. The plates were then incubated for 1-2 hours in the incubator, and the absorbance was measured at 450 nm using a microplate reader. Each experimental condition was performed in triplicate.

Cell viability of patient-derived lung organoids was evaluated using the CellTiter-Lumi™ Luminescent Cell Viability Assay. Organoids with mEGFR-driven NSCLC were seeded in 384-well plates, with approximately 2000 cells per well encapsulated in a matrix droplet. The following day, treatment was initiated with a range of cisplatin concentrations, starting at 100 µM and decreasing in four-fold dilutions as indicated, combined with 5 µM of the TBK1 inhibitor GSK8612. After 72 hours, cell viability was assessed by adding 25 µL of CellTiter-Lumi™ reagent to each well, followed by a 2-minute orbital shake to promote cell lysis and a 10-minute incubation at room temperature to stabilize the luminescent signal before measurement using a multi-mode microplate reader equipped with chemiluminescence detection capabilities.

### Caspase3/7 green probe apoptosis detection

Organoids were cultured on Matrigel in 24-well plates until they reached 50-60 µm in size to assess apoptosis in patient-derived lung organoids harboring mEGFR mutations. Organoids were then harvested by gentle suspension in pre-cooled PBS and seeded into 96-well plates at approximately 1x10^4 cells per well in 100 µL of lung cancer culture medium (Aime Medical, 10-100-010). Caspase-3/7 Green ReadyProbes™ Reagent (ThermoFisher) was added at 2 drops per mL of medium. Treatment included cisplatin at a maximum concentration of 100 µM and a 4-fold dilution series, alone or in combination with 5 µM of the TBK1 inhibitor GSK8612. Apoptotic activity was monitored over 72 hours using the Incucyte® S3 Live-Cell Analysis System (Sartorius) to capture and analyze changes in green fluorescence indicative of caspase-3/7 activity.

### Colony formation assay

Colony formation assays were performed to evaluate the clonogenic capacity of non-small cell lung cancer (NSCLC) cell lines H1975 and H1299. Cells expressing inducible control shRNA or shTBK1 were seeded at a density of 3,000 cells per well in 6-well plates. Following seeding, cells were treated with designated drugs and/or doxycycline. The culture medium, inclusive of drugs, was replenished every three days throughout 8 to 12 days to maintain optimal growth conditions. For observation, the colonies were fixed with methanol and stained with 0.2% crystal violet.

### Histopathological analysis

For histological examination, mouse lung tissues were dissected, fixed in 4% paraformaldehyde at 4°C for 12 hours, and dehydrated in a graded ethanol series. Following dehydration, tissues were embedded in paraffin and sectioned at 10 µm thickness. Deparaffinization was achieved by xylene treatment, and rehydration was performed through descending ethanol concentrations to water. Sections were then stained with Hematoxylin and Eosin (H&E) for general histological examination. Antigen retrieval was performed using EDTA buffer with microwave heating for immunohistochemical analysis. Sections were permeabilized with 0.5% Triton-X100 and blocked with 3% BSA to reduce non-specific binding. Overnight incubation at 4°C with primary antibodies (pEGFR, CST, 1:500) was followed by applying secondary antibodies and visualization with DAB staining. Processed sections were imaged using an automated pathology scanner.

### Nano-liquid chromatography/Tandem MS (Nano LC-MS/MS) analysis

Phoenix National Proteomics Core services performed Nano LC/tandem MS analysis for protein identification, characterization, and label-free quantification. Tryptic peptides were separated on a C18 column and analyzed by LTQ-Orbitrap Velos(Thermo). Proteins were identified using the National Center for Biotechnology Information search engine against the human or mouse RefSeq protein databases. The mass spectrometry proteomics data of TBK1 or STING modifications by mEGFR have been deposited to the ProteomeXchange Consortium (http://proteomecentral.proteomexchange.org) via the iProX partner repository with the dataset identifier PXD033926.

### RNA isolation and sequencing library preparation

Total RNA was isolated from samples using TRIzol (Thermo Fisher, 15596018), following the manufacturer’s protocol. The quantity and purity of the extracted RNA were assessed using a NanoDrop ND-1000 spectrophotometer (NanoDrop, Wilmington, DE, USA), and RNA integrity was evaluated on a Bioanalyzer 2100 (Agilent, CA, USA). Samples with concentrations >50 ng/μL, RIN values >7.0, and total RNA >1 μg were used for downstream analysis. Poly(A)-tailed mRNA was captured using two rounds of oligo(dT) magnetic beads (Dynabeads Oligo (dT), Thermo Fisher, 25-61005).

For library preparation, captured mRNA was fragmented at 94°C for 5 minutes in a fragmentation buffer (NEBNext® Magnesium RNA Fragmentation Module, NEB, E6150S). First-strand cDNA was synthesized using SuperScript™ II Reverse Transcriptase (Invitrogen, 1896649), followed by second-strand synthesis with E. coli DNA polymerase I (NEB, M0209) and RNase H (NEB, M0297). dUTP (Thermo Fisher, R0133) was incorporated into the second-strand synthesis to maintain strand specificity. Blunt-end DNA fragments were prepared, A-tailed, and ligated to T-tailed adaptors. After size selection and purification with magnetic beads, libraries were amplified using 14 cycles of PCR, targeting a fragment size of 300 bp ± 50 bp. According to standard procedures, libraries were sequenced in paired-end mode on an Illumina NovaSeq™ 6000 platform (LC Bio-Technology Co., Ltd., Hangzhou, China). Raw reads were processed and mapped to the human genome (hg19) using TopHat v2.1.1, and gene expression was quantified in fragments per kilobase of exon per million mapped fragments (FPKM) using Cufflinks v2.2.1. Genes with FPKM <1 in all samples were excluded from further analysis.

### Orthotopic tumor growth in nude mice

The nude mice were purchased from SLAC Laboratory Animal. H1975 cells (2 × 10⁶ viable cells) were resuspended in 50 μL of fresh RPMI 1640 medium and mixed with 50 μL of YEASEN Matrix. This cell suspension was then subcutaneously injected into six- to eight-week-old nude mice. Doxycycline (Dox) was administered in drinking water at a 1 mg/mL concentration at the indicated time points to induce the control shRNA or shTBK1 expression in nude mice. For chemotherapy and TBK1 inhibition, cisplatin (2.5 mg/kg body weight) was administered intraperitoneally every two days starting on day 3, while the TBK1 inhibitor GSK8612 (5 mg/kg body weight) was administered intragastrically (i.g.) daily starting on day 3. Tumor growth was monitored daily starting on day 7 or 8 post-injection, following protocols approved by the Institutional Animal Care and Use Committee (IACUC) of Zhejiang University. Tumor size is presented as a square caliper measurement and was calculated based on two perpendicular diameters(mm²). By ethical guidelines, mice were euthanized when the tumor diameter reached 15 mm and recorded as deceased due to tumor burden.

### In vivo study design

Rosa26^mEGFR^ transgenic mice with a C57BL/6J background were generated. Specifically, the SApolyA-CAG-LSL-EGFR(L858R-T790M)-IRES-luc-2A-tdTomato-WPRE-polyA expression cassette was precisely inserted into the Rosa26 locus via homologous recombination. Both male and female littermates were used for all the experiments. Adeno-CMV-Cre or Lenti-CRISPR-V2-Cre virus was delivered intratracheally to induce tumor formation in transgenic mice. Lungs were collected for analysis at the indicated time points. One month after the Cre-virus infection, cisplatin (2.5 mg/kg body weight) was administered intraperitoneally, and the TBK1 inhibitor GSK8612 (5 mg/kg body weight) was administered intragastrically, either alone or in combination. All the mice were bred and maintained in a pathogen-free animal facility at the laboratory animal center of Zhejiang University. The care of experimental animals was approved by the Committee of Zhejiang University and followed Zhejiang University guidelines.

### Statistics and reproducibility

Quantitative data are presented as the mean ± standard error of the mean (SEM) from at least three independent experiments. When appropriate, statistically significant differences between multiple comparisons were analyzed using the one-way or two-way ANOVA test with Bonferroni correction. Differences were considered significant at p<0.05. If preserved and properly processed, all samples were included in the analyses, and no samples or animals were excluded with conventional injection damage. No statistical method was used to predetermine the sample size. All experiments except those involving animals were not randomized. Immunoblotting, reporter assays, and qRT-PCR experiments have been independently repeated a minimum of three times to ensure reproducibility. The investigators were not blinded to allocation during experiments and outcome assessment.

## SUPPLEMENTAL FIGURE LEGENDS

**Supplemental Figure 1.**
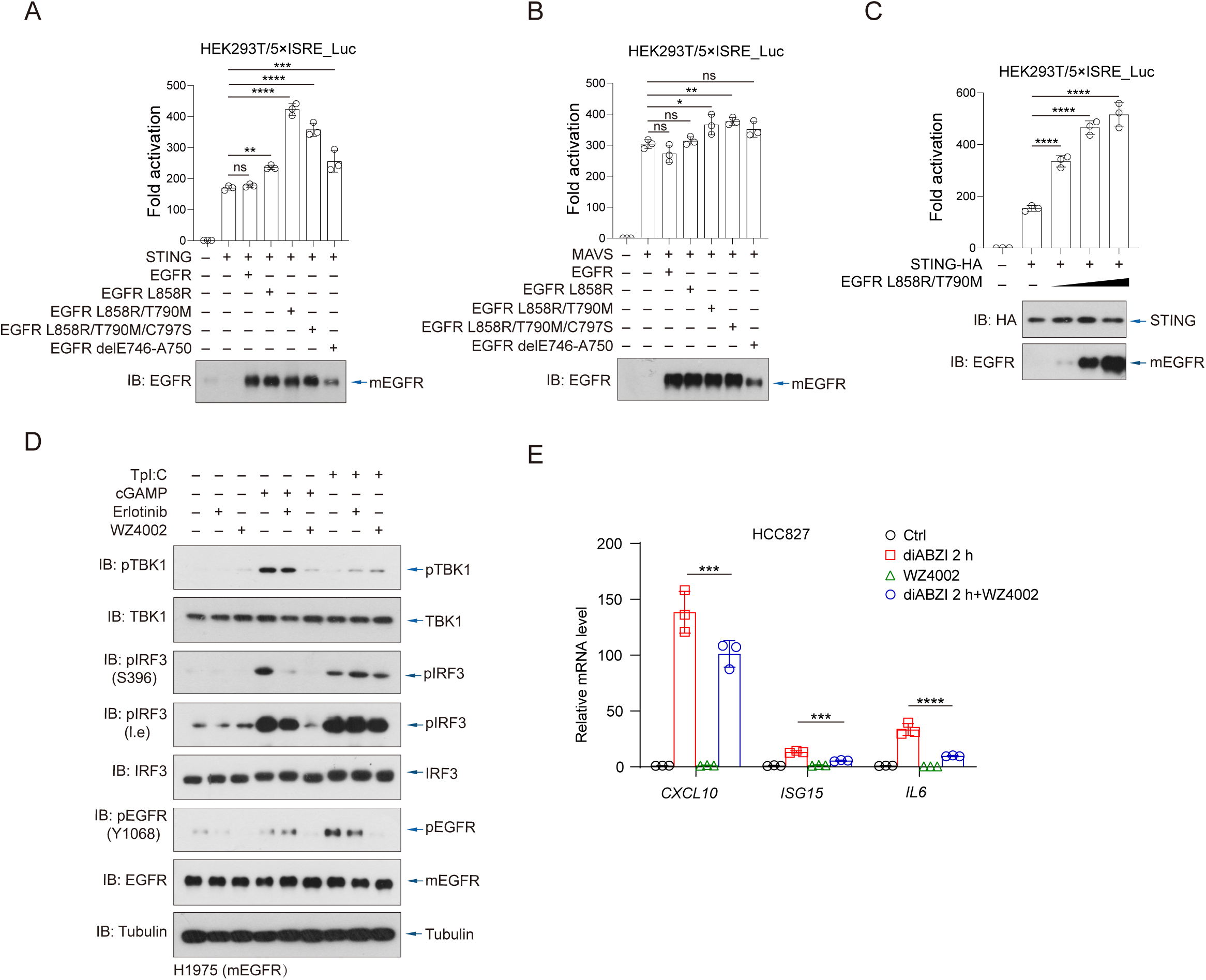
EGFR hotspot mutations kinase-dependently facilitate cGAS-STING signaling. **(A)** HEK293T cells were co-transfected with an IRF3-responsive ISRE-luciferase reporter, STING, and various EGFR constructs (L858R, L858R/T790M, L858R/T790M/C797S, or delE746-A750). Luciferase activity was measured 24 hours post-transfection and presented as fold induction relative to the empty vector control. EGFR and mutant EGFR expression were confirmed by immunoblotting. **(B)** HEK293T ISRE-Luc cells were co-transfected with MAVS and the indicated EGFR mutants to evaluate their effect on MAVS-mediated IRF3 activation. Luciferase activity is shown as fold induction compared to empty vector control. EGFR and mutant EGFR expression were validated by immunoblotting. **(C)** HEK293T ISRE-Luc cells were co-transfected with STING-HA and increasing amounts of EGFR L858R/T790M. Luciferase activity was quantified as a fold change relative to vector control. STING and EGFR expression were confirmed by immunoblotting. **(D)** H1975 NSCLC cells were treated with the STING agonist cGAMP (1 h), the dsRNA analog poly(I:C) (4 h), or vehicle, in the presence or absence of erlotinib or WZ4002. Phosphorylation and total levels of TBK1, IRF3, and EGFR were assessed by immunoblotting. **(E)** HCC827 cells were treated with diABZI (2 h), WZ4002, or the combination. mRNA levels of interferon-stimulated genes (*CXCL10*, *ISG15*, and *IL6*) were measured by quantitative RT-PCR. Data are presented as mean ± SEM (n = 3) and shown relative to untreated control cells. Statistical analysis was performed using one-way ANOVA with Bonferroni correction. ***, P<0.001, ****P < 0.0001.

**Supplemental Figure 2.**
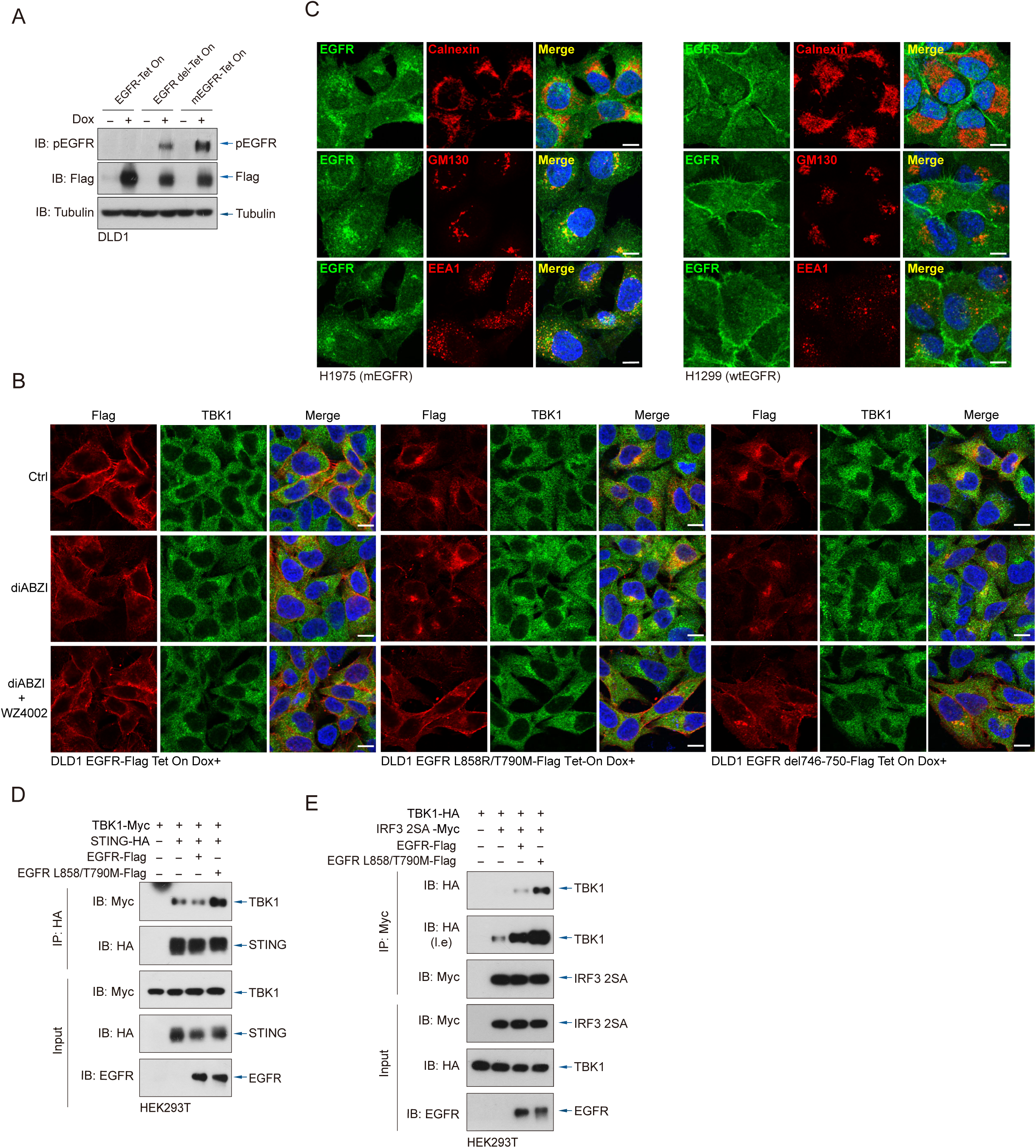
Mutant and wild-type EGFR differentially localize and interact with TBK1-STING signalosomes. **(A)** DLD1 Tet-On cells stably expressing Flag-tagged wild-type or mutant EGFR (L858R/T790M or delE746-A750) were induced with doxycycline (48 h). EGFR expression and phosphorylation were analyzed by immunoblotting. **(B)** DLD1 cells expressing the indicated EGFR variants were treated with diABZI (1 h) with or without WZ4002 (10 μM,6 h), then fixed and stained with antibodies against Flag (red) and TBK1 (green). Immunofluorescence images showed potential colocalization of EGFR and TBK1. Scale bars, 10 μm. **(C)** H1975 and H1299 cells were fixed and co-stained with antibodies against EGFR (green) and one of the following organelle markers, including calnexin (ER), GM130 (Golgi), or EEA1 (early endosomes) in red. Colocalization was evaluated by confocal microscopy. Scale bars, 10 μm. **(D)** HEK293T cells were co-transfected with Flag-tagged wild-type EGFR or EGFR L858R/T790M, HA-tagged STING, and Myc-tagged TBK1. C Lysates were immunoprecipitated with anti-HA and analyzed by immunoblotting with anti-Myc, anti-HA, and anti-EGFR antibodies. **(E)** HEK293T cells were co-transfected with Flag-tagged wild-type EGFR or EGFR L858R/T790M, Myc-tagged IRF3 2SA mutant (S385A/S386A), and HA-tagged TBK1. Myc immunoprecipitates were analyzed by immunoblotting.

**Supplemental Figure 3.**
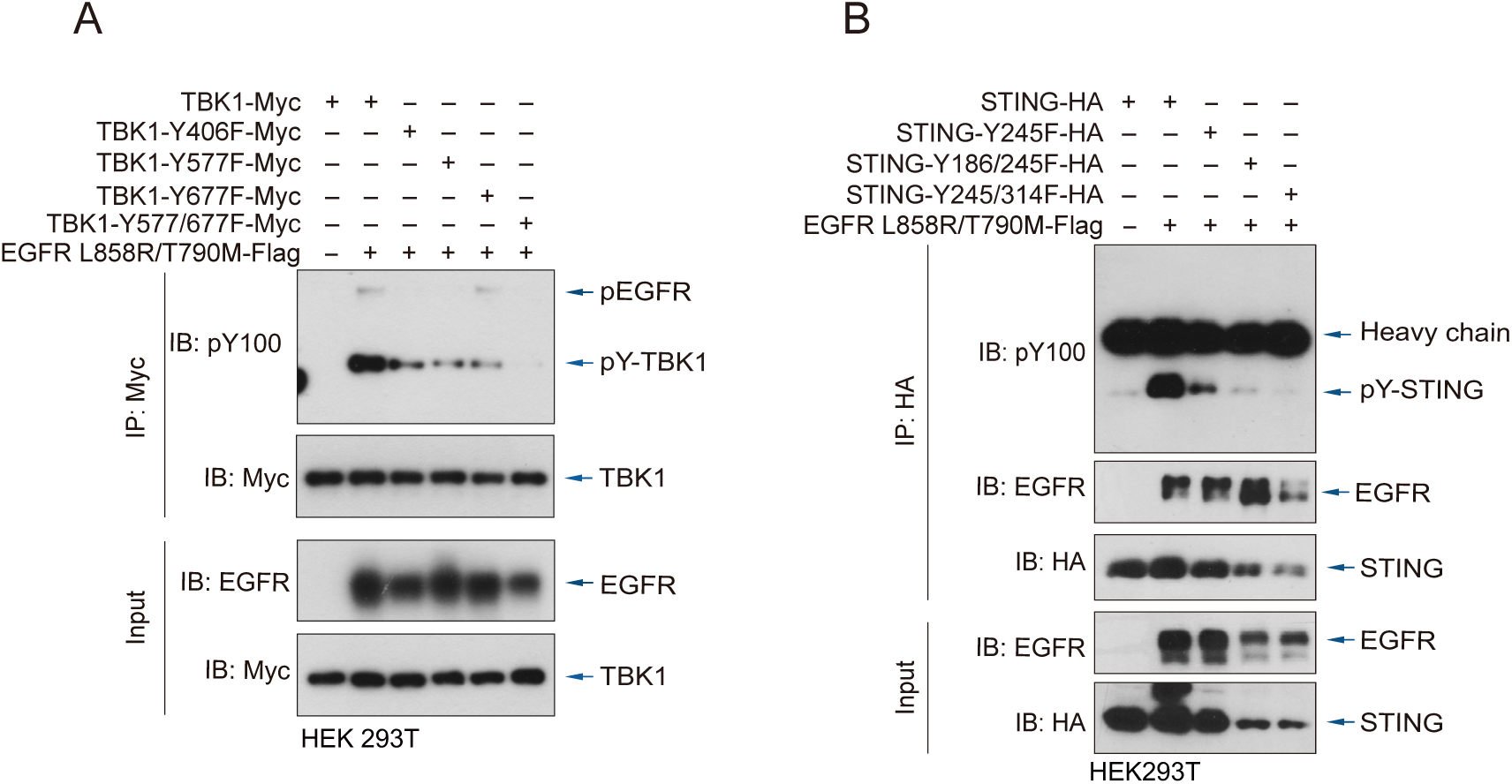
mEGFR tyrosine phosphorylates and facilitates STING and TBK1 activation. **(A)** HEK293T cells were co-transfected with EGFR L858R/T790M and either wild-type TBK1 or TBK1 mutants (Y406F, Y577F, Y677F, or Y577F/Y677F). After immunoprecipitation by anti-Myc, tyrosine phosphorylation was assessed using anti-pY100 immunoblotting. Mutation of individual tyrosine reduced EGFR-induced TBK1 phosphorylation to varying degrees, with the Y577F/Y677F double mutation showing the most significant reduction. **(B)** HEK293T cells were co-transfected with EGFR L858R/T790M and either wild-type STING or point-mutated forms of STING (Y245F, Y186F/Y245F, or Y245F/Y314F). HA immunoprecipitants were analyzed by immunoblotting with anti-pY100 to detect STING tyrosine phosphorylation.

**Supplemental Figure 4.**
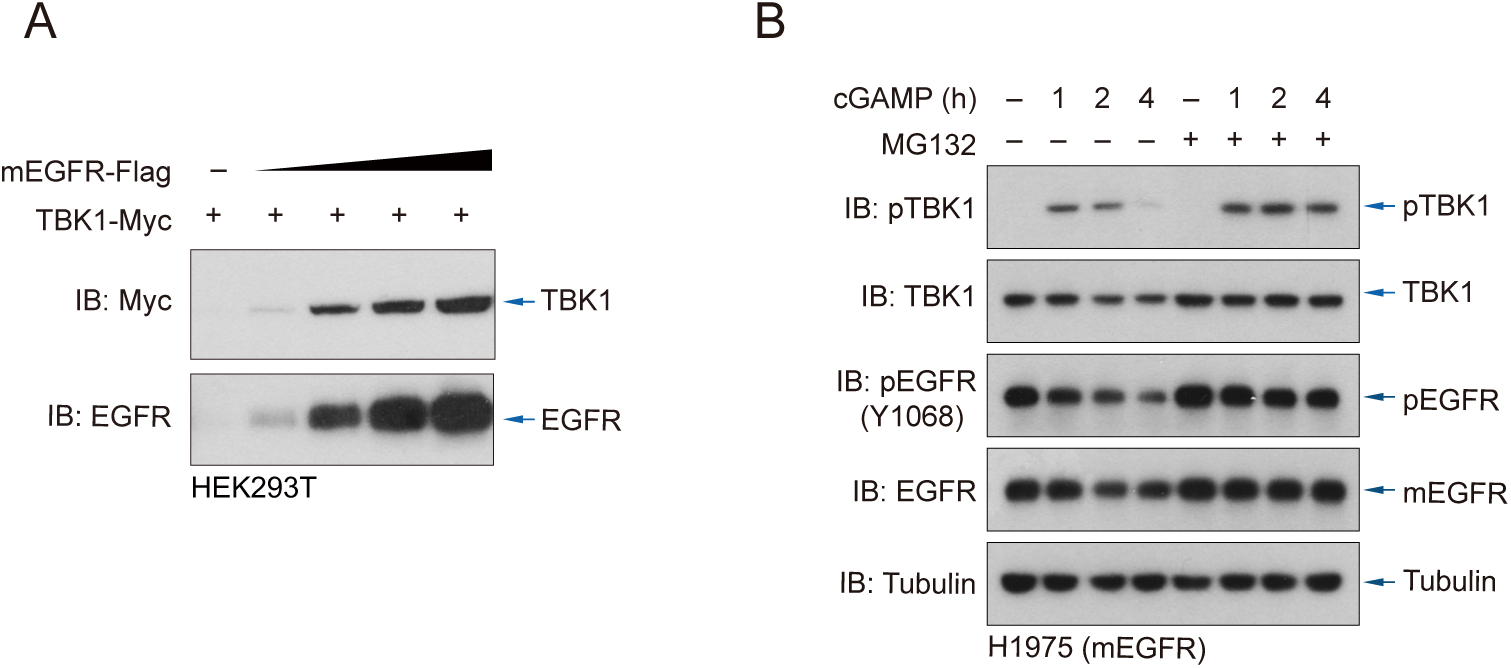
mEGFR-mediated phosphorylation hyperactivates and stabilizes TBK1. **(A)** HEK293T cells were co-transfected with TBK1-Myc and increasing amounts of Flag-tagged EGFR L858R/T790M. Immunoblotting revealed a dose-dependent increase in TBK1 protein levels corresponding to mEGFR expression. **(B)** H1975 cells were treated with cGAMP for the indicated time points in the absence or presence of the proteasome inhibitor MG132 (10 μM, 6 h). TBK1 protein levels were analyzed by immunoblotting to assess the impact of proteasomal inhibition on TBK1 stability.

**Supplemental Figure 5.**
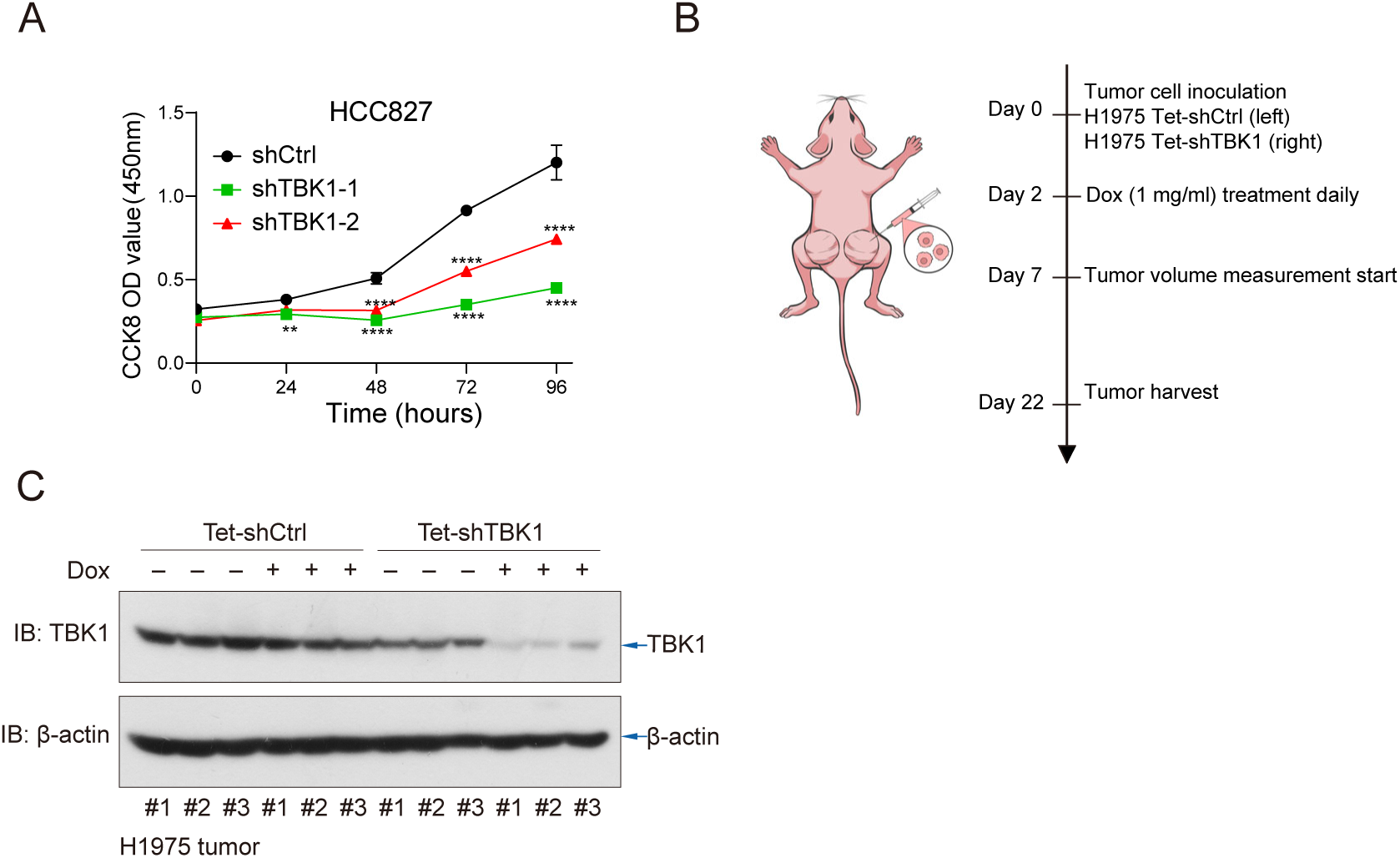
STING and TBK1 are indispensable for mEGFR-driven NSCLC survival. **(A)** CCK-8 assays measured cell viability in mutant EGFR-driven HCC827 NSCLC cells with shCtrl or shTBK1. Data are presented as mean ± SEM. **(B)** Schematic illustration of the in vivo experimental timeline for H1975 xenograft implantation and doxycycline treatment. **(C)** Immunoblotting analysis of TBK1 protein levels in individual H1975 tumor samples (n = 3 per group) with or without doxycycline treatment to induce shTBK1 expression.

**Supplemental Figure 6.**
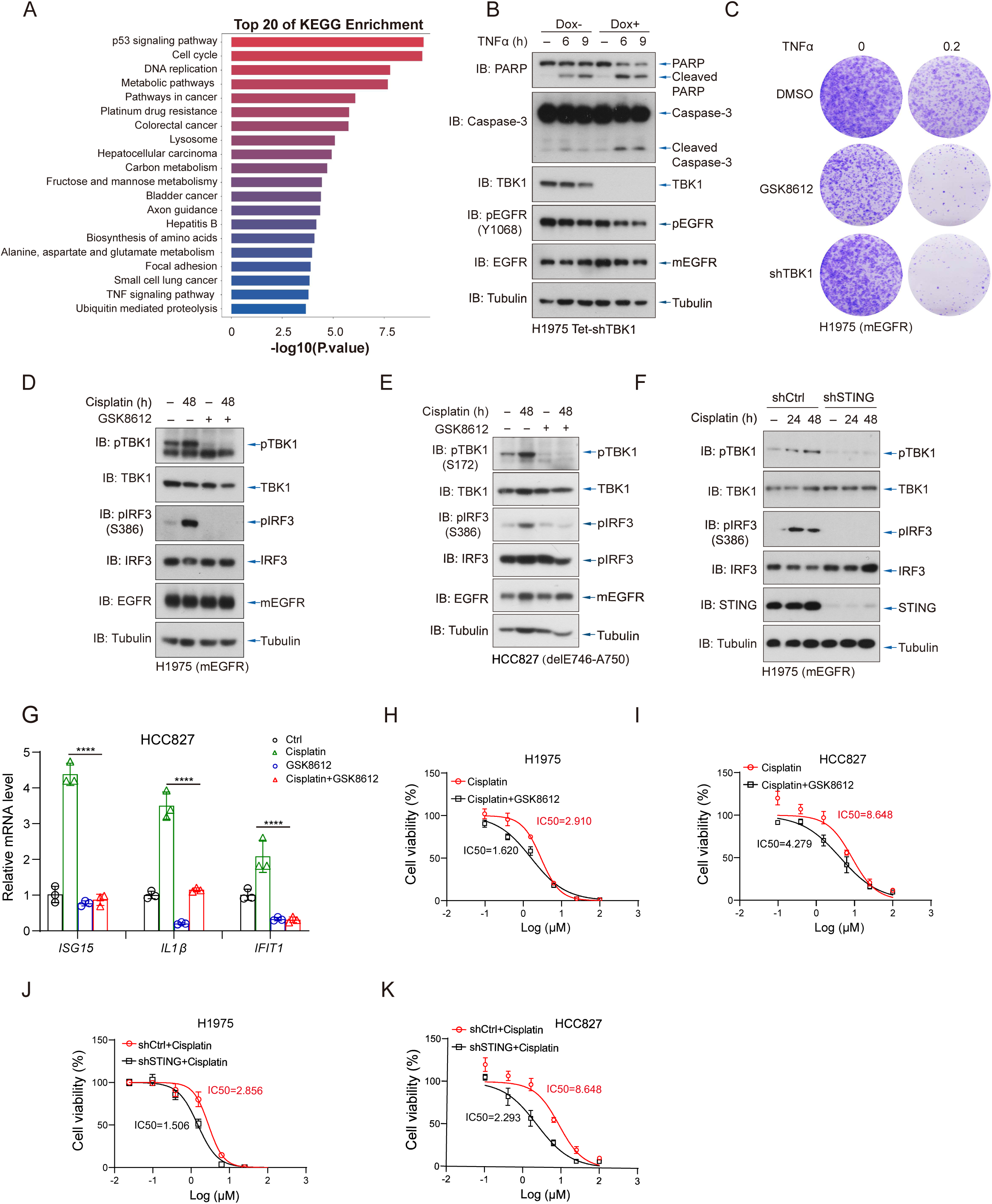
TBK1 facilitates DNA damage repair programs to survive mEGFR-driven NSCLC. **(A)** The transcriptome analyses of KEGG-enriched pathways for H1975 cells with TBK1 depletion indicated suppression of oncogenic and survival cellular processes. Y-axis: enriched pathways; X-axis: enrichment scores; color intensity denotes p-value significance. **(B)** Immunoblotting of H1975 Tet-shTBK1 cells treated with TNFα in the presence or absence of Dox for the indicated time points. TBK1 depletion exacerbated TNFα-induced apoptosis, evidenced by increased cleaved-PARP and cleaved-caspase-3 levels. **(C)** Colony formation assays in H1975 cells treated with TNFα and DMSO, GSK8612, or TBK1 shRNA. Both pharmacological inhibition and genetic depletion of TBK1 sensitized cells to TNFα-induced growth suppression. **(D–E)** Immunoblotting of H1975 (D) and HCC827 (E) cells treated with cisplatin (8 μM, 48 h), with or without GSK8612 (5 μM), to assess TBK1-IRF3 pathway activation. **(F)** Immunoblotting analysis of H1975 cells expressing shCtrl or shSTING and treated with cisplatin (8 μM) for the indicated time points, showing STING-dependent activation of TBK1 signaling upon cisplatin treatment. **(G)** Quantitative RT-PCR analysis of interferon-stimulated genes (*ISG15*, *IL1β*, and *IFIT1*) in HCC827 cells treated with cisplatin (8 μM), GSK8612 (5 μM), or the combination. TBK1 inhibition suppressed cisplatin-induced ISG expression. **(H–I)** Dose-response curves of cisplatin alone or combined with GSK8612 in H1975 (I) and HCC827 (J) cells. Co-treatment significantly decreased cisplatin IC_50_, suggesting enhanced sensitivity. **(J–K)** Dose-response curves of cisplatin in H1975 (J) and HCC827 (K) cells expressing control or STING-targeting shRNAs. STING depletion significantly reduced IC_50_ values of cisplatin, indicating increased sensitivity. Results in G and H–K are shown as mean ± SEM. Statistical analysis by ANOVA with Bonferroni correction; *P < 0.05, **P < 0.01, ***P < 0.001.

**Supplemental Figure 7.**
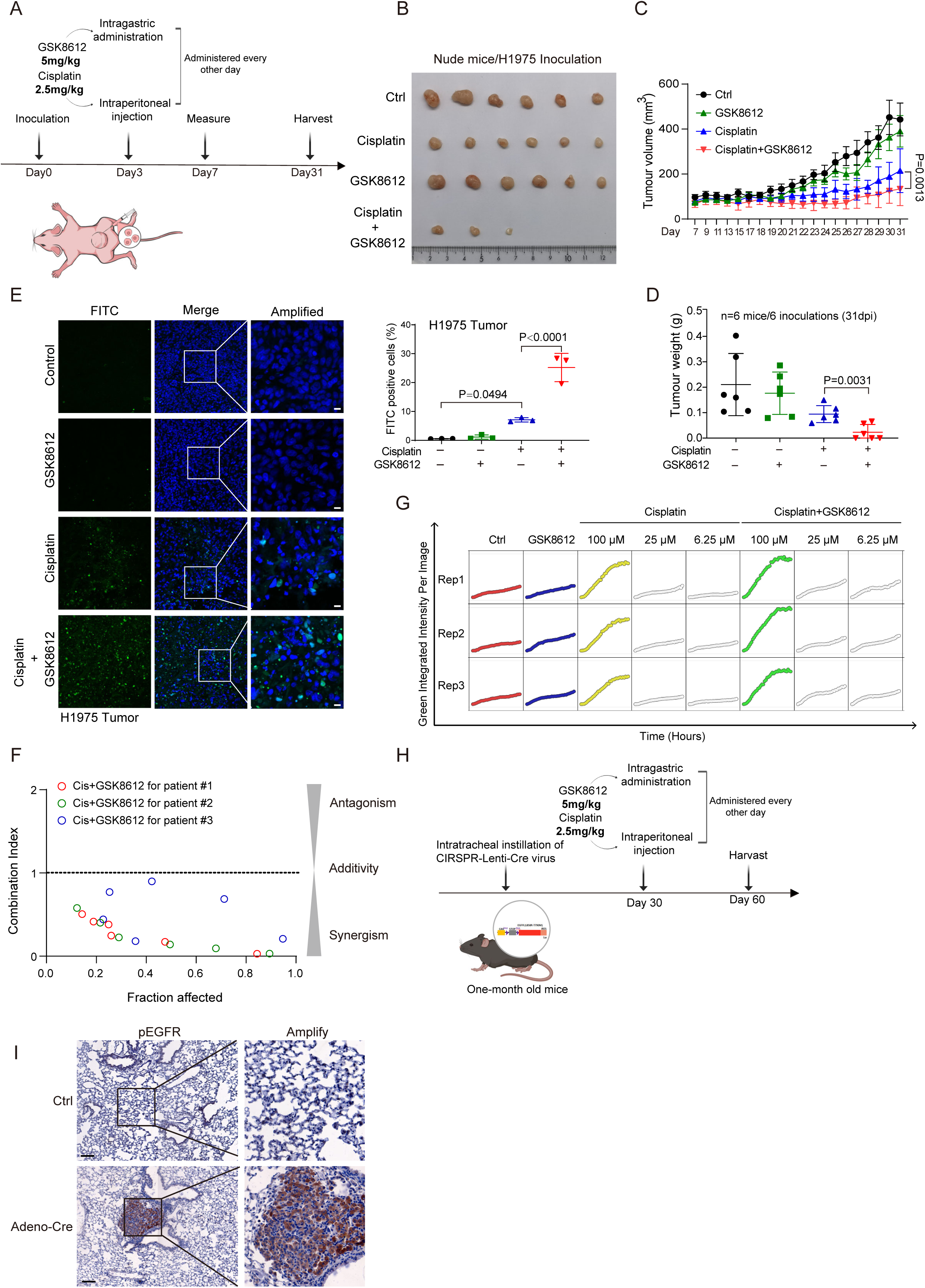
Combining cisplatin and TBK1i eradicates mEGFR-driven NSCLC PDOs and spontaneous tumors. **(A)** Schematic of the experimental design for the H1975 xenograft model. Nude mice were subcutaneously inoculated with H1975 NSCLC cells and treated every other day with GSK8612 (5 mg/kg, intragastric) and/or cisplatin (2.5 mg/kg, intraperitoneal), starting on day 3 post-inoculation. **(B)** Representative images of excised tumors at day 31 post-inoculation (dpi). Mice receiving combination therapy (cisplatin + GSK8612) exhibited the most significant tumor suppression. **(C–D)** Quantification of tumor progression in xenografts. Tumor volume was measured at the indicated time points (C), and tumor weight was assessed at 31 dpi (D). Combination treatment significantly reduced both parameters compared to monotherapies. Data are presented as mean ± SEM (n = 6). **(E)** TUNEL staining of xenograft tumor tissues showed markedly increased apoptosis in the combination group, as indicated by elevated FITC-positive cells (green). Scale bars, 10 µm. Quantification shown on the right (n = 3). **(F)** Drug interaction analysis using CompuSyn software demonstrated synergism (combination index, CI < 1) between cisplatin and GSK8612 across a range of concentrations in EGFR-mutant NSCLC PDOs from three patients. **(G)** Quantification of caspase-3/7 activation over time in PDOs (Patient #1) treated with cisplatin (at indicated concentrations) and GSK8612 (5 μM). Data are shown from three biological replicates using a live-cell apoptosis detection assay. **(H)** Schematic of the treatment schedule in the spontaneous mEGFR NSCLC mouse model. One-month-old mice were intratracheally instilled with CRISPR-Lenti-Cre virus to induce lung-specific expression of EGFR L858R/T790M. On day 30, mice received GSK8612 and/or cisplatin every other day until sacrifice on day 60. **(I)** Immunohistochemistry (IHC) of lung tissues from control or Adeno-Cre–infected mice revealed strong pEGFR expression in tumor regions of the Cre-treated group. Scale bars, 100 µm.

